# Medieval social landscape through the genetic history of Cambridgeshire before and after the Black Death

**DOI:** 10.1101/2023.03.03.531048

**Authors:** Ruoyun Hui, Christiana L. Scheib, Eugenia D’Atanasio, Sarah A. Inskip, Craig Cessford, Simone A. Biagini, Anthony W. Wohns, Muhammad Q.A. Ali, Samuel J. Griffith, Anu Solnik, Helja Niinemäe, Xiangyu Jack Ge, Alice K. Rose, Owyn Beneker, Tamsin C. O’Connell, John E. Robb, Toomas Kivisild

**Affiliations:** Alan Turing Institute; London, UK; McDonald Institute for Archaeological Research, University of Cambridge; Cambridge, UK; Estonian Biocentre, Institute of Genomics, University of Tartu; Tartu, Estonia; St John’s College, University of Cambridge; Cambridge, UK; Institute of Molecular Biology and Pathology, CNR; Rome, Italy; School of Archaeology and Ancient History, University of Leicester; Leicester, UK; Cambridge Archaeological Unit, Department of Archaeology, University of Cambridge; Cambridge, UK; Department of Human Genetics, KU Leuven; Leuven, Belgium; School of Medicine, Stanford University; Stanford, USA; Department of Genetics and Biology, Stanford University; Stanford, USA; Core Facility, Institute of Genomics, University of Tartu; Tartu, Estonia; Wellcome Genome Campus, Wellcome Sanger Institute; Hinxton, UK; Department of Archaeology, University of Durham; Durham, UK; Department of Archaeology, University of Cambridge; Cambridge, UK

## Abstract

The extent of the devastation of the Black Death pandemic (1346-53) on European populations is known from documentary sources and its bacterial source illuminated by studies of ancient pathogen DNA. What has remained less understood is the effect of the pandemic on human mobility and genetic diversity at local scale in the context of the social stratification of medieval communities. Here we study 275 newly reported ancient genomes from later medieval and post-medieval Cambridgeshire, from individuals buried before, during, and after the Black Death. The majority of individuals examined had local genetic ancestries. Consistent with the function of the institutions, we found a lack of close relatives among the friars and the inmates of the hospital in contrast to their abundance in general urban and rural parish communities. Accounting for the genetic component for height accentuates the disparities between social groups in stature estimated from long bones, as a proxy for health and the quality of life. While we detect long-term shifts in local genetic ancestry in Cambridgeshire that either pre- or postdate the Black Death, we find no evidence of major changes in genetic ancestry nor, in contrast to recent claims, higher differentiation of immune loci between cohorts living before and after the Black Death.

## Introduction

Evidence from ancient DNA (aDNA) continues to increase our understanding of the human past. By linking the genetic profiles to a place and time, it allows us to study population movements (Orlando et al., 2021; Skoglund & Mathieson, 2018), genetic relatedness (Cassidy et al., 2020; Kuhn et al., 2018; Ringbauer et al., 2021), infectious diseases (Spyrou et al., 2019; Warinner et al., 2017), and natural selection (Irving-Pease et al., 2021; Marciniak & Perry, 2017) as they occurred. When combined with historical and archaeological contexts, such information offers a more detailed perspective on life in past societies.

aDNA studies centered around broad geographical regions and long time periods have been fundamental in establishing major migration events, population turnovers or continuity in both prehistory and historic periods, while being less informative about everyday life experience within complex societies. Taking what we call ‘the whole town approach’, we have studied hundreds of skeletal remains from later medieval (*c*. 1000–1550 CE) Cambridgeshire (Table 1). They were excavated from burial grounds connected with different social groups: urban and rural parish churchyards, urban charitable institutions, and religious institutions. For historical context, we also included post-medieval burial grounds (*c*. 1550–1855 CE). Apart from Clopton and Hemingford Grey, all the sites are within a few kilometers of each other (Figure 1A).

**Table 1.**
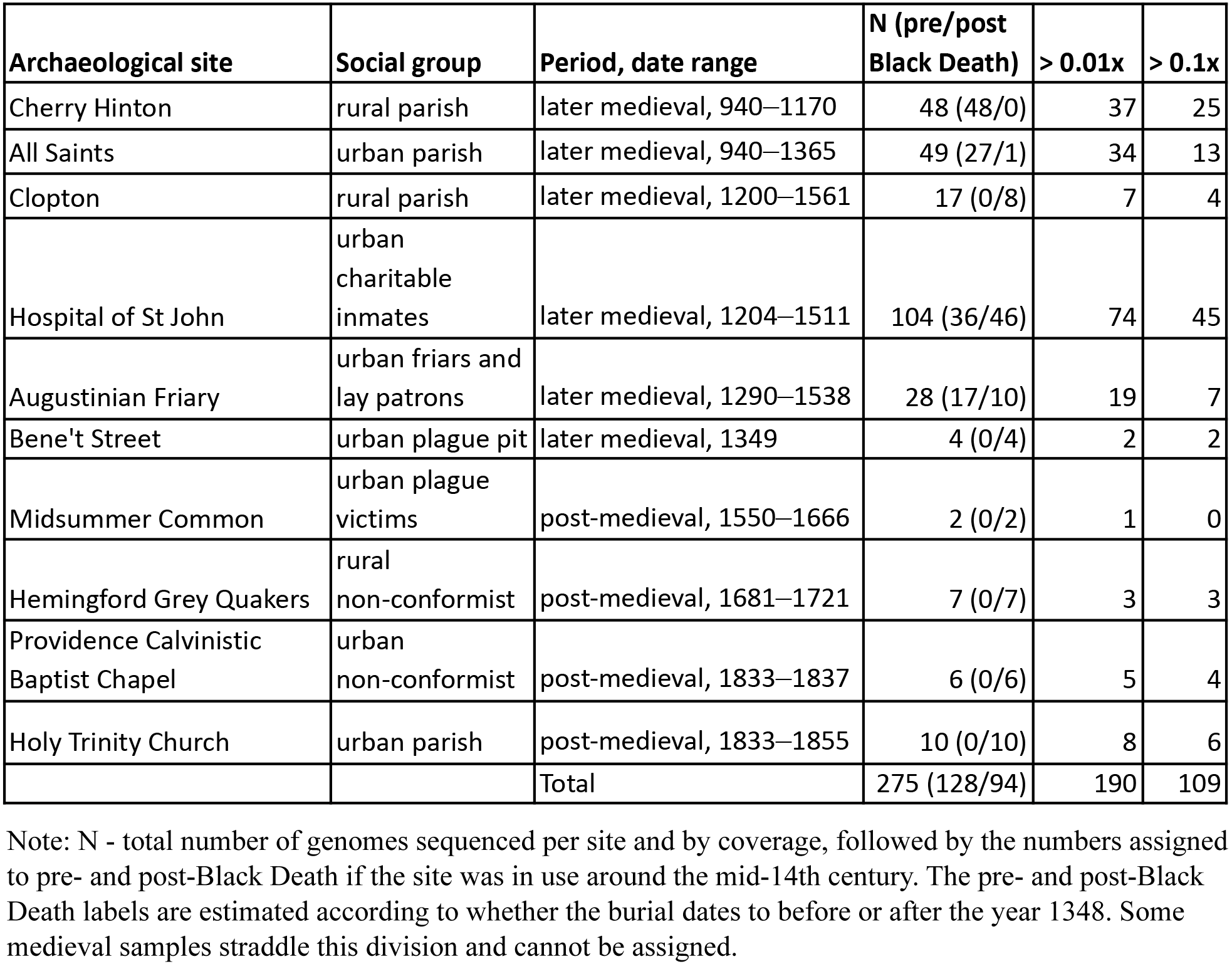
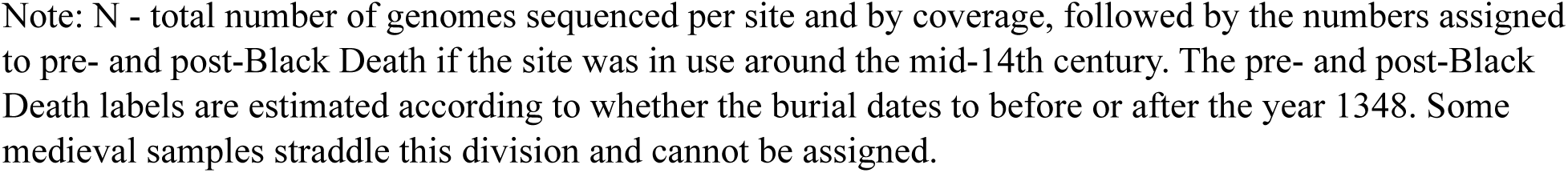
Ancient genomes from later medieval and post-medieval Cambridgeshire.

| Archaeological site | Social group | Period, date range | N (pre/post Black Death) | > 0.01x | > 0.1x |
| --- | --- | --- | --- | --- | --- |
| Cherry Hinton | rural parish | later medieval, 940–1170 | 48 (48/0) | 37 | 25 |
| All Saints | urban parish | later medieval, 940–1365 | 49 (27/1) | 34 | 13 |
| Clopton | rural parish | later medieval, 1200–1561 | 17 (0/8) | 7 | 4 |
| Hospital of St John | urban charitable inmates | later medieval, 1204–1511 | 104 (36/46) | 74 | 45 |
| Augustinian Friary | urban friars and lay patrons | later medieval, 1290–1538 | 28 (17/10) | 19 | 7 |
| Bene't Street | urban plague pit | later medieval, 1349 | 4 (0/4) | 2 | 2 |
| Midsummer Common | urban plague victims | post-medieval, 1550–1666 | 2 (0/2) | 1 | 0 |
| Hemingford Grey Quakers | rural non-conformist | post-medieval, 1681–1721 | 7 (0/7) | 3 | 3 |
| Providence Calvinistic Baptist Chapel | urban non-conformist | post-medieval, 1833–1837 | 6 (0/6) | 5 | 4 |
| Holy Trinity Church | urban parish | post-medieval, 1833–1855 | 10 (0/10) | 8 | 6 |
|  |  | Total | 275 (128/94) | 190 | 109 |

Note: N - total number of genomes sequenced per site and by coverage, followed by the numbers assigned to pre- and post-Black Death if the site was in use around the mid-14th century. The pre- and post-Black Death labels are estimated according to whether the burial dates to before or after the year 1348. Some medieval samples straddle this division and cannot be assigned.

**Figure 1.**
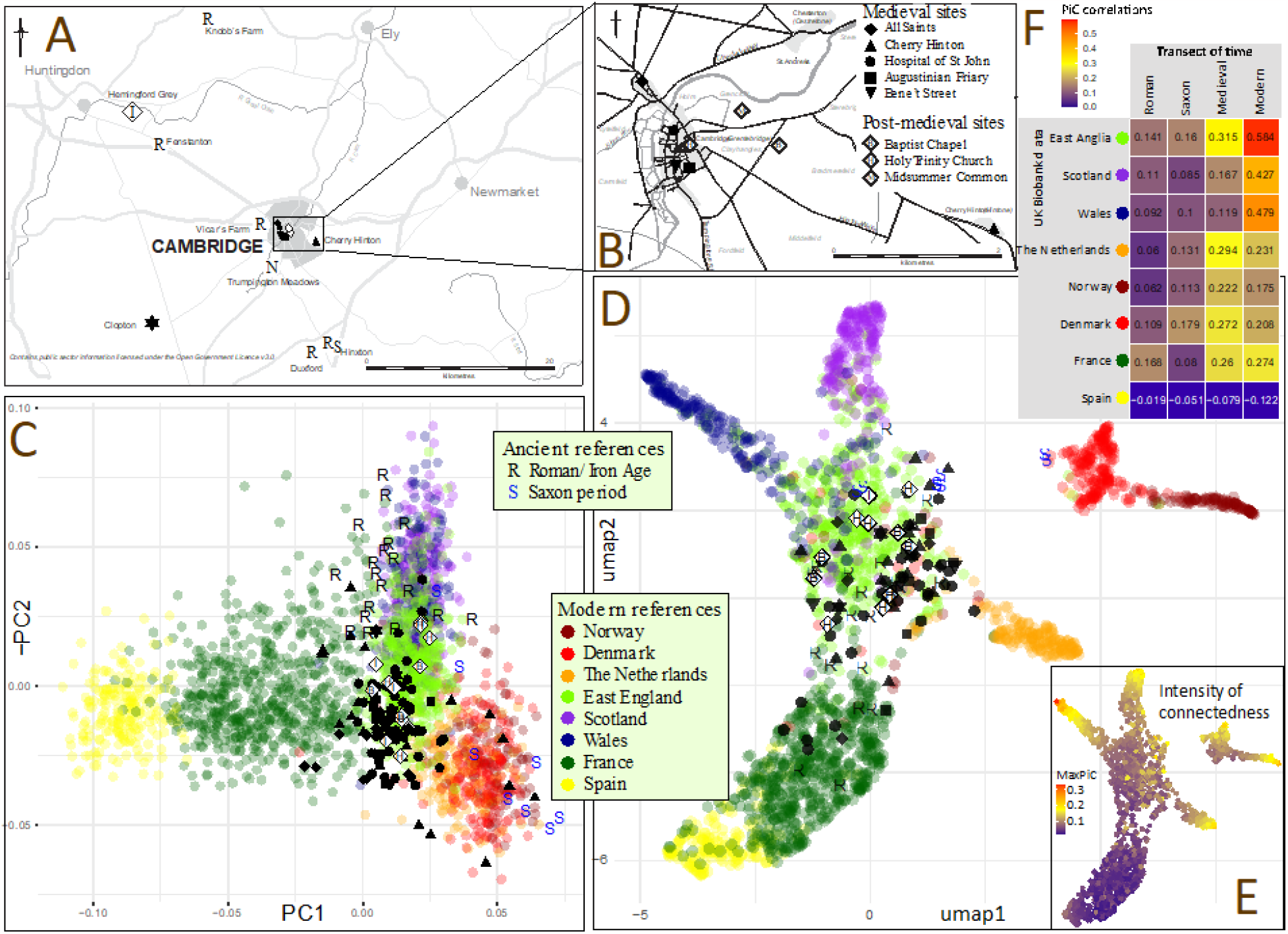
**A**. Sampling locations in Cambridgeshire, B zoomed in map of Cambridge C. PC plot of later medieval and post-medieval genomes with coverage > 0.1x in context of ancient and modern (UK Biobank) references, D. Supervised UMAP cluster-analysis using probability of individual connectedness (PiC) with modern references based on 5cM LSAI sharing among modern and ancient genomes with coverage > 0.2x, E. intensity of maximum PiC scores, F. Transect of time of correlations between the regional PiC vectors. The “Modern” correlation for East Anglia is shown as the correlation between PiC vectors of East Anglia and Bedfordshire/Hertfordshire.

Cambridge during the later medieval period was a middle-sized market town where people from all sections of society crossed paths. After they died, most townspeople were buried in one of the parish cemeteries, including All Saints by the Castle; however, other places of burial existed and increased over time. Towards the end of the 12th century, the Hospital of St John the Evangelist was founded by the townspeople as a charitable institute for the poor, the infirm and the sick. Most of the burials included in this study come from a cemetery for charitable inmates of the Hospital. The 13th century saw the founding of the university and houses of the mendicant orders, including an Augustinian Friary (from which some of our samples are drawn). Apart from the friars, some patrons of the friary were also buried in the cemetery and chapter house of the friary. The first wave of the Second Plague Pandemic (which we hereafter refer to with the commonly used term ‘Black Death’, although the term was not in use until the 18th century) hit Cambridge in 1349; some victims of it were found in a mass burial of unknown size on Bene’t Street (Cessford et al., 2021). Two parish cemeteries outside Cambridge, Cherry Hinton and Clopton, are within the rural hinterland of the town. Table 1 and Table S1 list the archaeological sites covered in this study, the dating of the burials, and the function of the medieval and post-medieval sites.

The Black Death and subsequent plague outbreaks had multiple effects on medieval society in England. Its death toll in Europe, estimated at 30%–65% (Benedictow, 2021; Horrox, 1995), could have posed selective pressure for better resistance to the plague. Genetic adaptations via the innate (Dumay et al., 2019) and adaptive immune system (Immel et al., 2021) have been proposed. Although reference bias can pose challenges to detect allele frequency changes due to natural selection (Gopalakrishnan et al 2022), a study of 206 aDNA extracts from individuals buried in London and Denmark before, during and after the Black Death revealed significant enrichment of immune genes among highly differentiated single nucleotide variants (SNVs) suggesting significant impact of the pandemic in shaping the disease susceptibility of surviving population (Klunk et al., 2022).

Besides genetic susceptibility, it is not clear to what extent social identity modified the morbid and mortal effects of the Black Death. Plague mortality appeared to be selective with respect to frailty (DeWitte & Wood, 2008) caused possibly by factors such as malnutrition, impaired immunocompetence and others affected by social conditions. For example, the Great Famine of 1315–1322 could have severely affected people of low socio-economic position. Short stature as an indicator of poor health possibly increased the chances of dying during the pandemic (DeWitte & Hughes-Morey, 2012). In this sense health inequality between social groups sets the background for understanding the potential for different experiences through the pandemic. As longer-term consequences of the mortality, it has been argued that the Black Death pandemic initiated or accelerated profound socioeconomic changes, such as increased social mobility, improved quality of life of the laboring population, and technological innovations to increase productivity (Herlihy & Cohn, 1997; Campbell, 2016). Together with evidence from osteology, isotopes and the rich context around the burial grounds, we aim to explore to what extent genetic data might aid the construction of a social history, both in relation to the pandemic and regarding the more stable aspects of later medieval life.

## Results

We extracted aDNA and generated whole-genome shotgun sequence data from a total of 250 later medieval and 25 post-medieval skeletons, retrieving for further analyses 190 genomes at coverage > 0.01x (Table 1, Table S1). They form the most extensive bioarchaeological sampling within a focused temporal and geographical range to date. The examined medieval sites represent burials of individuals from different social and cause of death backgrounds, including urban cemeteries of the charitable poor from the Hospital of St. John, All Saints parish cemetery, Augustinian Friary, Bene’t Street plague burial and rural cemeteries of Cherry Hinton and Clopton (Table 1, Figure 1A, Supplementary Material). The analyses of the medieval genomes were performed in context of post-medieval genomes from four sites in Cambridgeshire (Table 1) as well as published genomic data of the Late Iron Age/Roman (*c*. 100 BCE–400 CE) and Early Saxon periods (*c*. 400–700 CE) from Cambridgeshire and elsewhere from England (Martiniano et al., 2016; Schiffels et al., 2016). Average endogenous human DNA content was 13% and average contamination rate 1.14%, with 231 samples under 5%. Average damage in the first 5 base pairs was 8.02% (Table S1). A subset of 143 genomes sequenced to > 0.05x coverage were imputed to study the changes in phenotypes related to health and lifestyle. The imputed genomes include 109 individuals with coverage > 0.1x, which were subsequently used to resolve genetic ancestry, kinship, recent inbreeding, and heterozygosity.

### Genetic ancestry

The frequency of mtDNA haplogroups in England has remained relatively stable since the Neolithic (Table S2). PCA reveals that all individuals we sampled from Cambridgeshire share their ancestry with modern northern and western European populations without evidence of migration from more distant regions (Figure 1C, Figures S1-11). In contrast to genomes of individuals whose remains date to the Roman or Early Saxon periods (Martiniano et al., 2016; Schiffels et al., 2016), most later medieval genomes cluster with those from the modern English individuals from the UK Biobank data (Figure 1C). Individual outliers who, similarly to the majority of Early Saxon period individuals, are placed among modern Dutch and Danish populations, include a few from Cherry Hinton (Figure S2) and the Hospital of St John (Figure S5). Two of them (PSN332 from the Hospital and PSN930 from Cherry Hinton) are also outliers in terms of dental enamel ^87^Sr/^86^Sr values (PSN332 = 0.7122, PSN930 = 0.7108), in comparison to the rest of the Cambridgeshire population sampled (Rose & O’Connell, forthcoming). These values, particularly for PSN332, are higher than the estimated biosphere ^87^Sr/^86^Sr values for the East of England (Evans et al., 2018), indicating that they did not spend their childhoods in the area local to where they were buried (Rose & O’Connell, forthcoming). Although some areas in Britain, particularly Wales and Scotland as well as Cornwall and smaller areas across England, do have estimated biosphere ^87^Sr/^86^Sr values which could produce the values in these individuals, combined with the genetic information, PSN332 is more likely to be first-generation migrant from the Scandinavian peninsula. This is consistent with an influx of continental northern European ancestry after the Roman period, followed by increased affinity to present-day Scandinavian populations since the Viking Age (Gretzinger et al., 2022). For instance, in this period there was an active North Sea trade network linking eastern England, Norway and northern Germany and this would be a plausible origin for PSN332. On the other hand, three individuals from Cherry Hinton, seven from All Saints, two from the Hospital, one from the Friary, and a post-medieval Midsummer Common burial clustered closely with modern French samples from the UK Biobank in the PCA result (Figure 1B, Figures S2-11).

To study the genetic affinity changes across time at finer geographic resolution we defined inter-individual connections by identifying long (> 5cM) shared allele intervals (LSAI-s) with IBIS (Seidman et al., 2020) and explored the modularity of individual connectedness, PiC (Kivisild et al., 2021), among the historical and modern genomes. Similarly to PCA results, we find that the majority of historical genomes from Cambridgeshire cluster by their connectedness with modern UK Biobank genomes from East England (Figure 1D, Table S2) whereas a small fraction of later medieval and Roman period genomes, which display low connectivity (Figure 1E), cluster with the UK Biobank donors born in France who display low connectivity among themselves. The Early Saxon period genomes show higher connectivity with Scandinavian genomes, which is also reflected in individual PCA outliers from Cherry Hinton. Overall we observe regional shifts in individual connectedness over time (Figure 1F). There is increasing Danish connectivity in the transition from Roman to Early Saxon period; later, during and after the later medieval period, there is an increase of connectivity with both modern Dutch genomes (mirroring documentary evidence showing the Dutch as the most common late medieval immigrants locally (Ormrod et al., 2019, 2020)), and genomes from a broader zone of England. Finally, we observe a major shift in modern East England towards higher connectivity with Wales and Scotland, clearly reflecting the political and economic integration of recent Britain. Our analyses of individual connectedness in the People of the British Isles (POBI) (Leslie et al., 2015) data suggest that all later medieval genomes from Cambridgeshire likely draw most of their genetic ancestry from the same sources as present-day central/eastern England population (Figure 2). Although we are able to distinguish certain regional differences in the modern data with our approach, such as between Cornwall and Devon or between North and South Yorkshire, we observe less resolution and lower connectedness, likely because of high mobility in the last few centuries, in a broad area between Lincolnshire and Surrey where our ancient genomes come from (Figure 2). This means that even if some of the individuals had come from Kent or Lincolnshire, for example, we would not be able to detect such fine-scale migration patterns among regions within that area.

**Figure 2.**
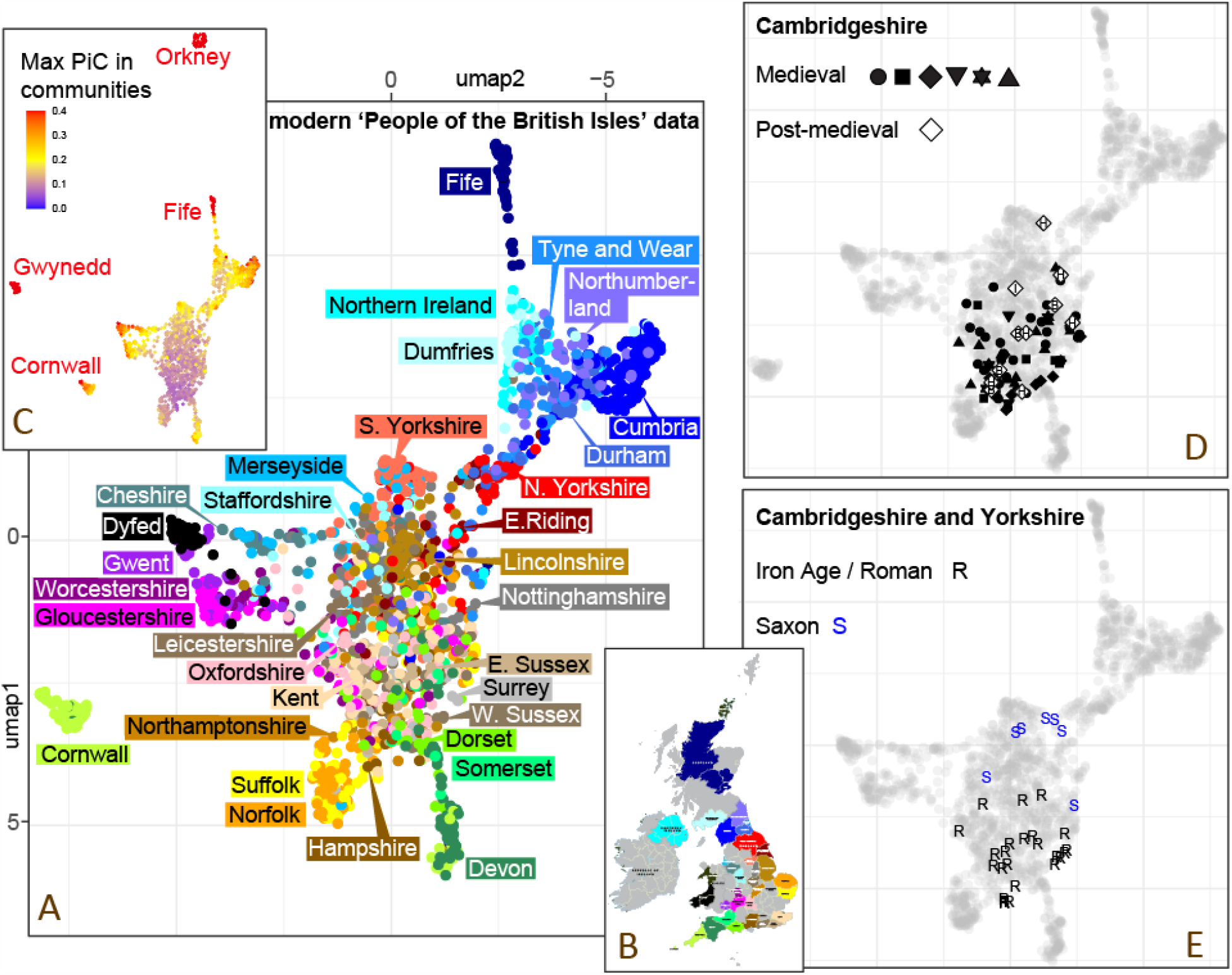
UMAP plot of individual connectedness among modern and ancient genomes from Britain. **A**. UMAP dimension reduction plot of individual connectedness among modern genomes of the ‘People of the British Isles’ project based on PiC scores of 20 significant communities with more than 10 members extracted from the combined data with the Louvain method (unsupervised cluster analysis). **B**. Map showing the color codes by counties for the modern genomes used in the UMAP plot A. **C**. Density of maximum PiC score values per individual in one of the extracted communities. **D**. UMAP coordinates of the medieval and postmedieval genomes (>0.2x coverage) from Cambridgeshire. Archaeological site codes as shown in Figure 1. **E**. UMAP coordinates of the Iron Age/Roman and Saxon period genomes.

Changes in genetic ancestry or selective pressure could cause phenotypic changes over time. We analyzed 214 individuals from the Roman period to the 19th century, including previously published data (Margaryan et al., 2019; Martiniano et al., 2016; Schiffels et al., 2016), for changes in allele frequency of 113 phenotype informative SNPs related to diet, health and pigmentation (Tables S4–S6). Out of 74 SNPs related to health and diet, only two involved in autoimmune disease reached the adjusted significance threshold in the ANOVA statistical tests showing differences between the medieval and post-medieval periods. Similarly, we cannot find significant changes during and after the later medieval period for the 39 SNPs affecting eye, hair and skin color included in the HIrisPlex-S set (Chaitanya et al., 2018) the few significant SNPs mainly driven by a change in the allele frequency that occurred during or just after the Roman/Iron Age period (Gretzinger et al., 2022; Martiniano et al., 2016).

### Social landscape

#### Kinship and relatedness

Although the “kinship” bonds that tie together social groups often go beyond or replace “blood-relationships”, the types and intensity of genetic relatedness among individuals buried in the same locality can be informative of the social structure of the population. To study the probability of genetic relatedness among burials of different social backgrounds we used READ-based estimates (Kuhn et al., 2018) of pairwise differences in autosomes and the X chromosome. Among individual pairs with > 10,000 overlapping SNPs, twenty-one cases of 1st–3rd degree relatedness were detected (Figure 3A, Table S7). All kinship pairs detected by READ that involved individuals with > 0.1x coverage were confirmed in our analyses with IBIS (Seidman et al., 2020) to share multiple > 7cM LSAI segments and kinship coefficient > 0.005 suggestive close degree of genetic relatedness (Table S7). Unsurprisingly, considering the time gaps, in analyses involving 463,855 UK Biobank samples none of the 97 later medieval samples showed kinship with modern samples when using 7cM threshold. Nine of the twelve tested post-medieval individuals are found to form a total of twenty 4th–6th degree relationships with modern individuals who identify themselves as British, including one born outside of the UK. Among the later medieval samples we detect 12 cases of more distant form of relatedness within the same archaeological site beyond those identified with READ - ten at Cherry Hinton and two at All Saints - while none found at the Hospital or Friary. We find multiple cases (more than 1% of all pairs considered) in the rural Cherry Hinton and urban All Saints parish cemeteries. In contrast, we detected only one pair of relatives, a middle-aged (46–59) friar and a female child (2nd degree). No kinship relations were found at the Hospital, despite the large sample size analyzed.

**Figure 3.**
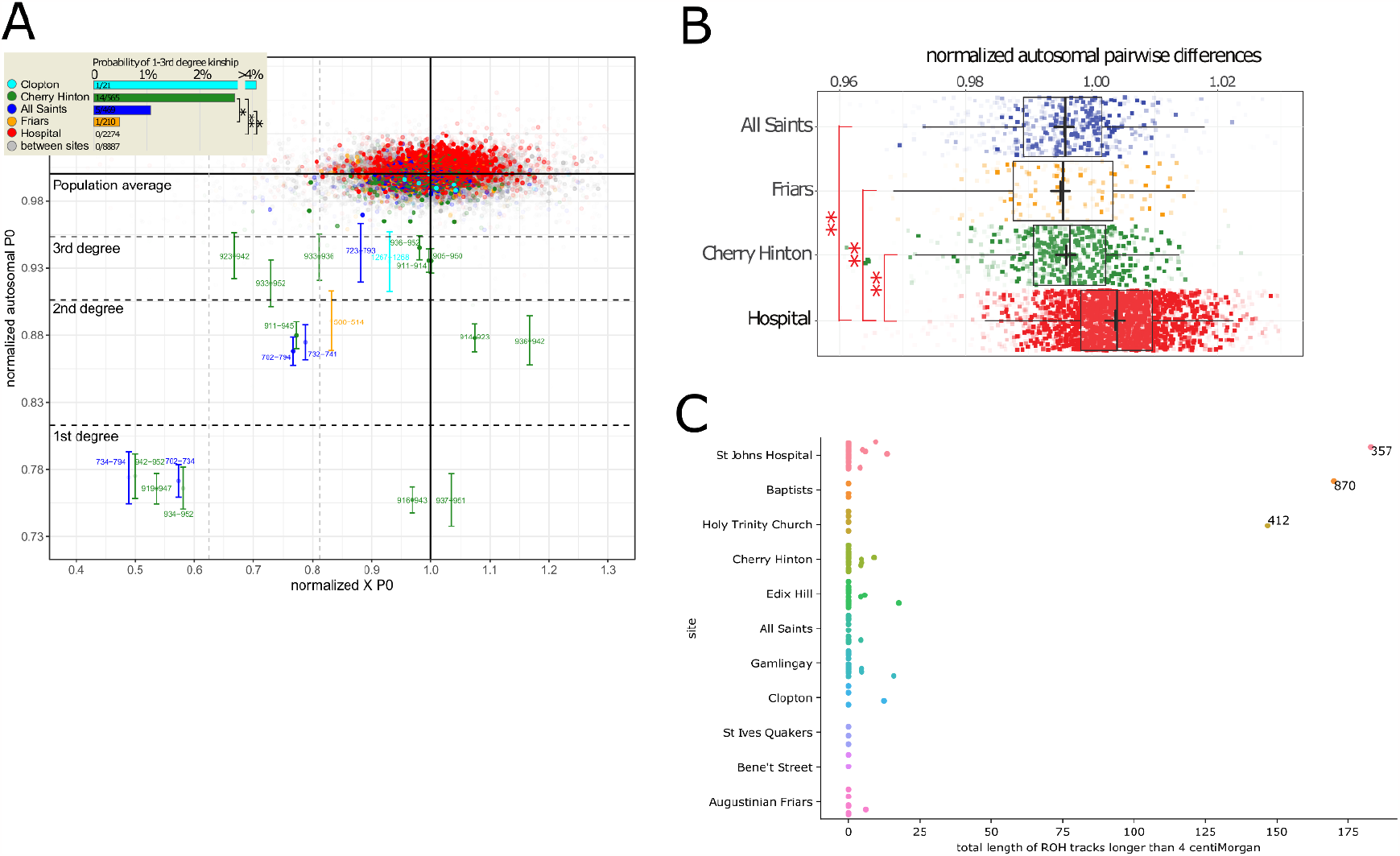
**A. Normalized pairwise differences (P0) in autosomal data and X chromosome for later medieval sites with more than five burials**. Each individual data point represents a pair of samples (from a total of 171 individuals with > 0.01x coverage tested), the aggregate coverage of which is reflected by the opacity of the color. Boundaries for the 1-3rd degree of relatedness for autosomal data were defined as in (Kuhn et al., 2018). Error bars with two standard errors of the mean are shown for the pairs of 1-3rd degree of relatedness only. ** and * correspond to significant differences at p < 0.01 and p < 0.05 by two-tailed t test, respectively. **B. Boxplot of normalized autosomal pairwise differences in four Cambridge medieval sites represented by the largest sample size in this study**. Each rectangular data point represents a pairwise comparison of individuals sampled from the same site, normalized (with READ) by the average of all pairwise comparisons made in the pool of all later medieval samples from Cambridge. The opacity of the rectangular color reflects the aggregate of the coverages of the two individuals. The results of significant (p < 0.01) two-tailed Wilcoxon rank sum test are shown with **. **C. The total lengths of ROH tracks longer than 4 centiMorgan in individuals grouped by site, showing highly inbred outliers**.

On average, pairwise differences between individuals from the Hospital were found to be greater than those between individuals from other sites (Figure 3B), highlighting the heterogeneity of the ancestry of individuals entering the Hospital. This finding reflects the way inmates were recruited into the Hospital, a charitable institution which took in individuals based on need, not residence or relatedness.

We also searched for runs of homozygosity (ROH) tracks in the imputed genomes using hapROH (Ringbauer et al., 2021) (Table S8). These are long segments where the maternal and the paternal copies of the genome are identical; a large amount of long ROH tracks would suggest recent inbreeding. Most individuals have no or very few ROH tracks that are longer than 4 centiMorgan, while three individuals (PSN357 from the Hospital, PSN870 from the Baptist Chapel and PSN412 from Holy Trinity) have up to 150–175 centiMorgan in ROH, which is compatible with the parents being 4th to 5th degree relatives, including 2nd to 3rd cousin marriage (Figure 3C).

#### Burden of health

Adult stature has been used as a proxy for health and quality of life in anthropological studies (Bogin, 2020; Vercellotti et al., 2014). Accounting for the genetic potential for height would better expose the effect of environmental stressors in people’s lives. Previous studies have shown that polygenic risk score (PRS) models developed in modern studies also have predictive power in ancient skeletal remains (Cox et al., 2022; Marciniak et al., 2022). We applied 8 PRS models for height (Chung et al., 2019) on our imputed medieval genomes, and compared the PRS with osteological stature estimated from long bone lengths (Buikstra & Ubelaker, 1994; White & Folkens, 2005). Most PRS models yield a correlation of 0.2–0.3 between the PRS and sex-adjusted stature estimation (Table S9). Different long bones would give slightly different stature estimation, among which the femur length has been shown to correlate most strongly with stature (White & Folkens, 2005). Accordingly, the correlation between the PRS and estimated stature increases when we limit to stature estimated from the femur only: for individuals whose genomes were sequenced to > 0.05x, the best-performing model (lasso using dosage data) explains 0.18 of the variation in stature estimated from all available long bones, increasing to 0.29 in stature estimated from the femur only. Similarly, when we increased the sequencing coverage cutoff for the genomes, the prediction accuracy increased from 0.18 at 0.05x (using all available long bones) to 0.28 at 0.2x. The highest accuracy we achieved is comparable to that in modern populations when the validation and training datasets come from separate studies (0.28 for the lasso model, compared to 0.39 when both come from the same study (Chung et al., 2019)), although we are likely to have benefited from a much larger imputation reference panel compared to the original study. We believe that the prediction accuracy of PRS will continue to improve if we further increase the sequencing depth.

As a balance between the sample size and the predictive power of PRS scores, we chose the PRS scores calculated from a lasso model and limited the samples to those with sequencing coverage above 0.05 with femur measurements (n = 88) for further modeling. We explored how the PRS scores and other variables, such as sex and social class, can predict stature using Bayesian linear regression models. Figure 3A illustrates the result of regressing stature on PRS scores and sex. The 95% credible interval of the posterior distribution of the coefficient for the PRS score is 0.20–0.40, showing that PRS models developed in modern populations can also explain a substantial amount of stature variation in the later medieval Cambridgeshire population.

When we limit the analysis to the core later medieval period (n = 76, excluding two prosperous benefactors buried in the Friary) and include social group as a predictor in addition to sex, adding PRS scores increases the model fit (difference in ELPD = 6.1, sd = 3.7) (Table S10). Moreover, including PRS scores as a predictor slightly increases the difference between sites (Figure 3B, Table S10): for example, 87% of the posterior distribution of the stature difference between charity cases and townsfolk is negative, suggesting people who ended up as inmates in the Hospital suffered more developmental stress; it increases to 90% after accounting for PRS score increases. We found similar trends regarding taller stature associated with the friars and shorter stature associated with the rural population.

### Before/After the Black Death

Since our pre/post-Black Death grouping comes from expert archaeological judgment over whether the burial occurred before or after the year 1348, some individuals in the post-Black Death group might have been born before 1348, although two obvious cases from Bene’t Street have been excluded from analysis. This might limit our power to detect changes after the Black Death. Our analyses of genetic ancestry (Figures 1-2, S1-11, Table S2) were unable to detect changes in rates of long distance migration associated with the Black Death comparable to those recently shown in case of Trondheim population (Gopalakrishnan et al., 2022). However, besides the potential effect on broader regional ancestry, the Black Death pandemic could have left other detectable signatures on genetic diversity of the population at genome-scale, or, it could have affected specific genes and variants associated with infectious disease vulnerability. To examine its effect on the genetic diversity of the Cambridge medieval population further, we estimated heterozygosity and nucleotide diversity genome-wide and in the HLA locus in imputed genomes.

Genome-wide heterozygosity and nucleotide diversity are sensitive to demographic events such as bottlenecks, founder events or admixture. High mortality during the pandemic could be detectable in extreme cases in a small isolated population as a reduction of diversity across all loci. Changes in the HLA region might capture possible signals of selection specifically at immunity loci. If any one or a few variants in this locus responded to selection they would have been expected to affect the whole region due to linkage. Consistent with the long-term effect of balancing selection on HLA locus, we find this region has higher density of heterozygous positions at common variants and nucleotide diversity in our pooled sample of imputed genomes (Figure 5A, Figure S13). However, the “before” and “after” the Black Death cohorts do not show higher than average allele frequency differentiation within the HLA region (Figure 5B) nor notable differences in the heterozygote density (Figure 5C). Within a subset of 50 imputed genomes assigned to either before or after the Black Death and coverage > 0.1x we observed no significant differences in genome-wide (2-tailed t-test, p = 0.205) nor HLA locus (p = 0.700) heterozygosities (Figure S12). Similarly, we did not detect changes in nucleotide diversity in the HLA region or genome-wide after the Black Death (Figure S13).

**Figure 4.**
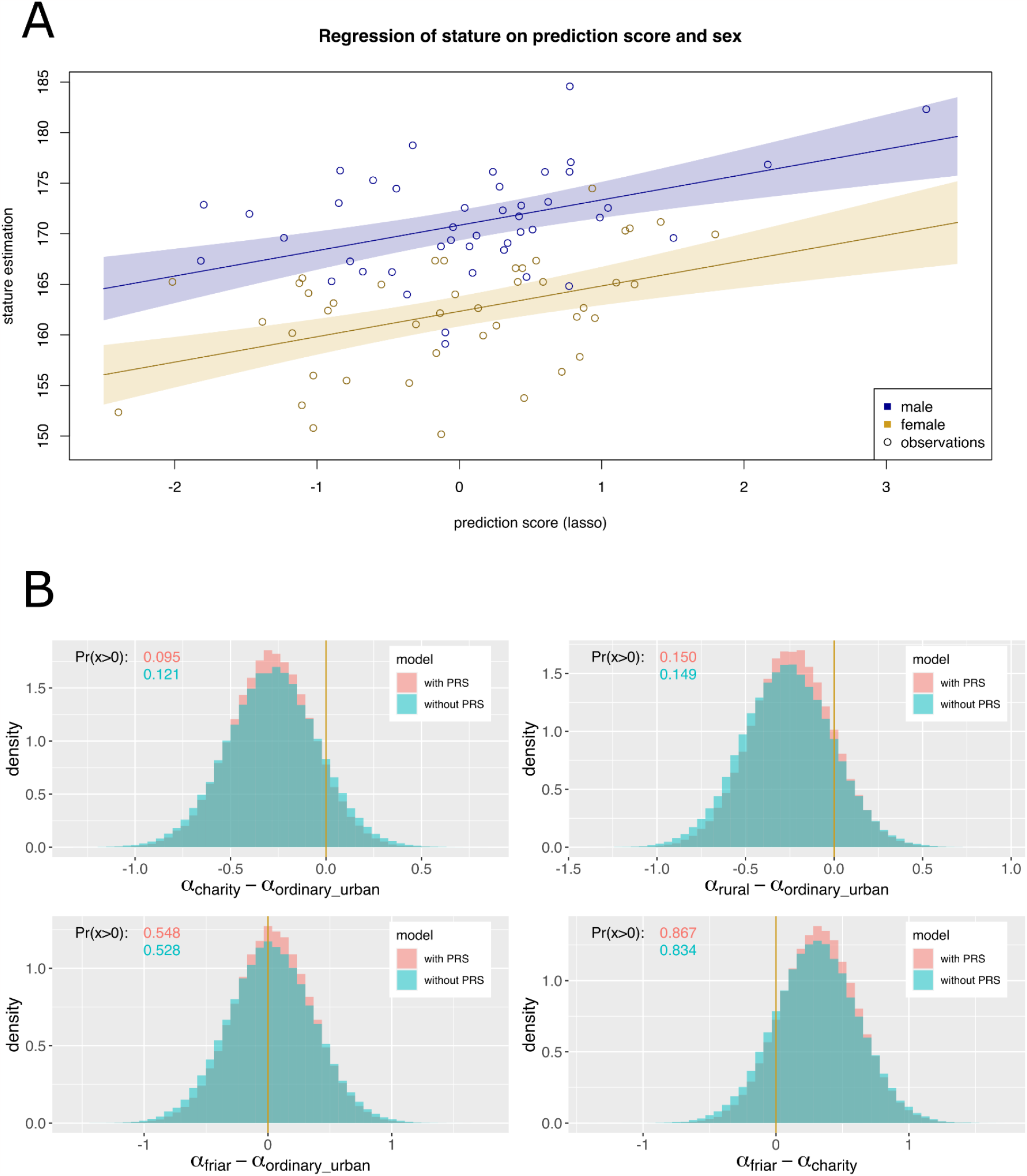
A: stature estimates from femur and PRS score for height in medieval individuals (coverage > 0.05x), showing regression lines with 90% posterior credible interval. B: posterior distributions of the difference in stature between social groups, with and without PRS score as a predictor in the linear model.

**Figure 5.**
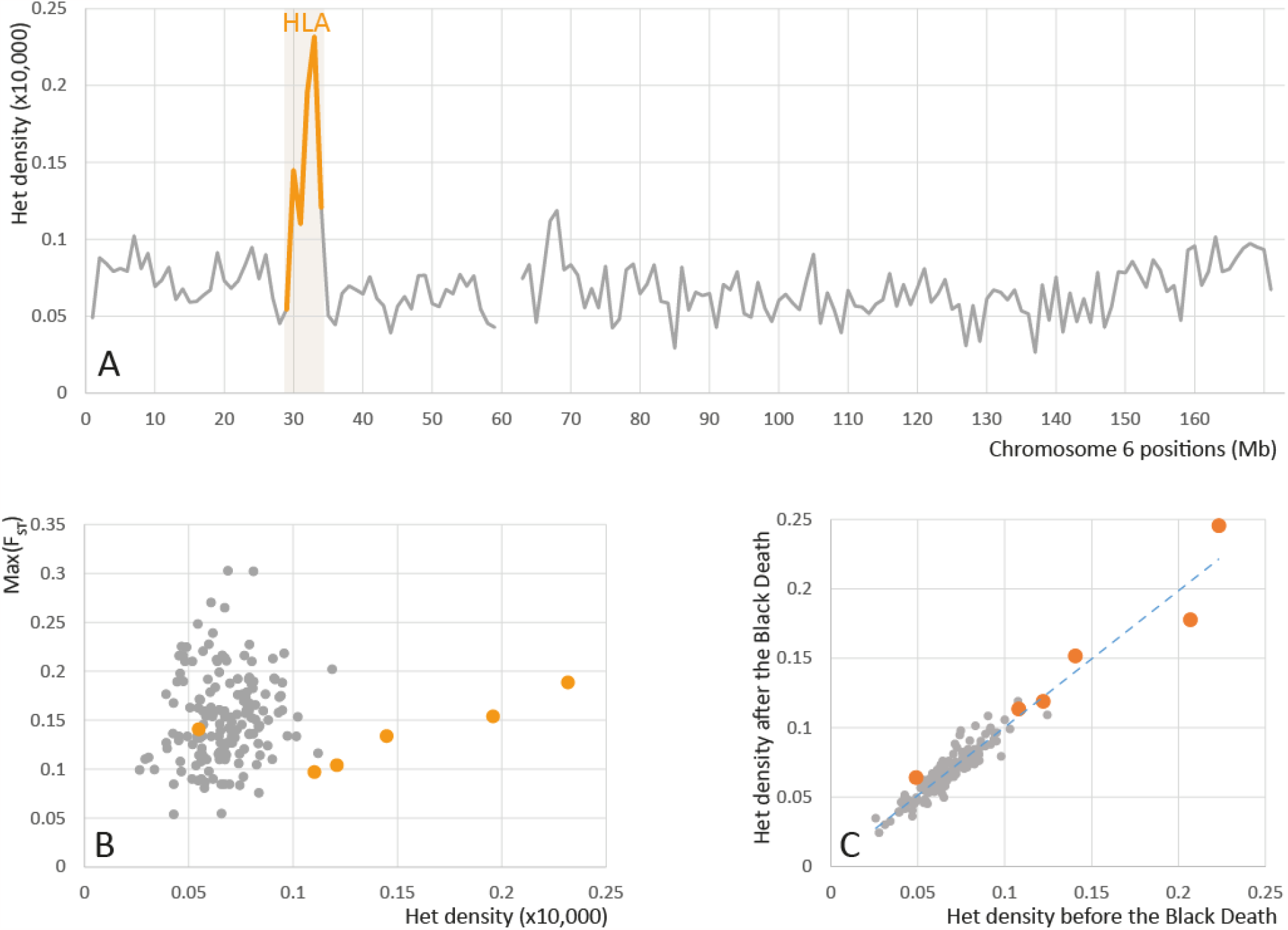
Heterozygote density and allele frequency differentiation in the HLA locus. **A**. Distribution of average heterozygote density in 1 Mbp windows of chromosome 6. Grey line shows the density of heterozygous sites at common variant positions with minor allele frequency higher than 0.05 in the HRC imputation panel in chromosome 6 for 50 imputed (> 0.1x coverage) pre- and post-Black Death genomes from Cambridge. The orange line highlights windows containing genes in the HLA locus. **B**. Scatter plot of Max(F_ST_) - maximum Fst between before and after the Black Death cohorts - and Het density values by 1 Mbp windows of chromosome 6. **C**. Het density in the before (n=31) and after (n=19) the Black Death cohorts.

Our analyses of 70 pre- and post-Black Death imputed genomes for changes in allele frequency in 25 variants previously identified as potential targets of selection in humans against viral and bacterial pathogens as well as four variants recently highlighted as selection targets specifically against *Y. pestis* (Klunk et al., 2022) revealed (Table S4, S6) one significant (p = 0.003) difference at individual test level at the rs42490 SNP in the *RIPK2* gene. The *RIPK2* allele previously shown to be protective against leprosy (Zhang et al., 2009) showed increased allele frequency after the Black Death. This result would not remain, however, significant after applying multiple test corrections and considering the limited sample size of our cohort it requires further validation in an independent data set.

None of the four immunity variants identified by Klunk et al. (Klunk et al., 2022) with significant allele frequency changes both in their London and Danish cohorts were replicated between Cambridge before and after Black Death cohorts (Table S11). We did observe a similar enrichment (1.4-fold, p = 0.0001) of variants related to immunity among highly differentiated variants (Fst > 95th percentile) when using the same list of immunity-related and neutrally evolving variants as the authors (Tables S11-12). Out of the 245 highly differentiated immunity-related variants identified in their London cohort, 22 were replicated, significantly more than expected by chance (p = 0.0001); However, 10 of the 22 overlapping variants that are above the 95th threshold in the Cambridge cohort and two of the three variants above the 99th threshold show opposite directionality of allele frequency change in time in London and Cambridge cohorts (Table S11). While the minor allele frequencies of the immunity variants appear to be highly correlated between our studies (r = 0.90, p < 1×10^−15^) the Fst between the pre- and post-pandemic cohorts are not (r = 0.019, p = 0.26). Importantly, the significant enrichment of immunity genes cannot be reproduced with our data when using the full list of 37,574 neutral regions defined by (Gronau et al., 2011) instead of the relatively small number of variants ascertained by Klunk et al. in its subset of 250 regions (Table S12). We observe a reduction (1.14 fold, p = 0.29) of high Fst values among the Klunk et al. immunity variants when we define the neutral 95th threshold using 55,965 variants from the full range of the 37,574 neutral regions, which becomes significant (1.73 fold, p < 1×10^−10^) when also using an expanded set of immunity variants from InnateDB (Table S12). Notably, within the pool of highly differentiated immune locus variants identified by Klunk et al. we observe significant excess of ‘gwas’ variants, i.e. positions that had previously been confirmed to be polymorphic (Table S12) over immunity variants ascertained in the exonic regions, suggesting that ascertainment of new variants from low coverage data (as by Klunk et al. in their ‘exon’ and ‘neutral’ categories) is one possible cause for the disappearance of the signal when we used the full set of neutral regions defined by Gronau et al. that overlap with positions confirmed to be polymorphic in the HRC panel to define the threshold.

## Discussion

Our analyses of genetic ancestry in Cambridge through the transect of time revealed, firstly, a notable shift in PCA and a corresponding increase of connectivity with Scandinavia between the Roman period and more recent samples. These findings likely reflect the cumulatively massive changes in local genetic ancestry in East England that occurred over many centuries, with 38% Saxon and 76% North Sea zone contributions estimated by previous studies (Gretzinger et al., 2022; Schiffels et al., 2016). Most of the individuals that fall outside of the local range of variation come from the rural cemetery at Cherry Hinton, possibly reflecting a diversity of Late Anglo-Saxon, Scandinavian and Norman ancestries in the process of intermixing. Despite the scale of these ancestry changes, it is difficult to convincingly detect any first-generation migrants from other parts of western Europe using genetic information alone, partially due to an absence of comparable reference material from medieval Europe. In case of the Friary, a context where individuals with non-local ancestry would have been more likely to be found, it is also possible that some migrating friars living in Cambridge during their life moved back and were buried elsewhere. Secondly, with the analyses of individual connectedness patterns we were able to distinguish between modern Dutch, Danish and Norwegian ancestries and observed a major increase of population-scale connectedness in Cambridge with the Dutch dating to the later medieval period. This pattern could in principle reflect gene flow either from the Low Countries to Cambridge or gene flow from East England to the Low Countries, although the former is better supported by documented sources. Finally, the major change we observed between medieval and modern periods in connectedness with modern Welsh and Scottish genomes is likely to reflect recent and ongoing migrations and mobility. In contrast to the highly region-specific LSAI sharing between modern and medieval genomes in Estonia (Kivisild et al., 2021), later medieval Cambridge genomes do not show increased affinity to modern East Anglia compared to other regions. These recent changes in genetic ancestry suggest that modern Biobank sources may not be ideal references for the study of population histories, highlighting the need for ancient DNA sampling of historic populations from different time points.

Although we observe major long-term changes in genetic ancestry, we did not observe a change in ancestry or genetic diversity between our pre- and post-Black Death cohorts in line with the hypothesized increased mobility after the Black Death due to labor shortages. The short time span of a few decades, small sample size and relatively low genetic differentiation in central and eastern England (Figure 2; (Leslie et al., 2015) would all make detecting short to medium-range migration within England challenging. The lack of evidence for long-distance migration is in agreement with earlier mtDNA evidence in medieval British and Danish cohorts (Klunk et al., 2019) but in contrast to genome-wide evidence of increased long-distance migration from medieval Trondheim (Gopalakrishnan et al., 2022). These results suggest that the pandemic had a variable impact on human mobility in European cities depending on their regional, socio-economic and political context and circumstances.

We were unable to detect differences in genetic ancestry between the ordinary urban population buried at All Saints, rural population buried at Cherry Hinton, charitable inmates buried at the Hospital of St John, and friars and benefactors buried at Augustinian Friary in the later medieval period, although historical record suggests that some friars might have traveled from as far as the Mediterranean to attend the Augustinian *studium generale* (regional study centre) here. The lack of close genetic kinship is consistent with the function of the Hospital as a safety net for those without family support, although we note that genetic relatedness does not capture the whole range of social relationships viewed as kinship. Relatedly, the hospital inmates also show elevated background genetic diversity compared to other later medieval sites. Some of them could have arrived from farther away than the average townsfolk, and the lack of local social support might be related to their hardship. Given that friaries maintained long-lasting ties with local benefactor families, it is not surprising that a female child who was a 2nd-degree relative to one of the friars was allowed to be buried within the Friary, but no other close genetic relatives were found among the Friary population.

We found a small number of cases of extended ROH tracks in the genome consistent with the parents being second to third cousins. These cases can be considered unexpected because marriage within four degrees of consanguinity was forbidden by the Church (Caserta, 2007). We speculate that this could result from incomplete church records, especially if the two parents attended different parish churches, or extra-pair paternity events in the family history.

We did not observe differentiation in genetic ancestry between the social groups, but we have shown that controlling for genetic information can help highlight variations caused by environmental stress. The predictive accuracy of the PRS scores for height in our later medieval samples is comparable to that seen in independent modern validation datasets (Chung et al., 2019) and remarkably higher than in previous studies using ancient genomes (Cox et al., 2022; Marciniak et al., 2022), possibly owing to 1. better PRS models; 2. a more genetically homogenous population, that is also 3. closer to the modern study population both in terms of genetic background and way of life, compared to the timescale of tens of thousands of years in the other studies. In our samples, controlling for PRS scores as an estimate of the genetic potential for height increases the difference between the effects attributed to social groups. The increase is limited, most likely because the social groups in this study do not differ much in their genetic potential for height. The transferability of PRS models developed on modern study population to historical population depends on population history, genetic architecture of the phenotype of interest, and environmental heterogeneity (Carlson et al., 2022); but we believe better genetic data and better understanding of complex traits will open up new exciting scope to disentangle the genetic effects and life experience in bioarchaeological studies.

The Black Death has been hypothesized to exert selective pressure on genes related to health and immunity (Immel et al., 2021; Stephens et al., 1998). If existing genetic variations differ in their resistance against the plague, and if the plague was a major cause of mortality, we would expect a drop in genetic diversity immediately after the pandemic. However, among individuals who lived shortly before or after the Black Death, we did not find changes in heterozygosity in the HLA region. Although the mortality rate (30-65%) was devastating, followed by subsequent plague outbreaks until the second half of the 17th century in Britain, it may not have exerted a strong enough uniform selective pressure for a long enough period to leave a detectable signature in the low-coverage genomes. One study has identified allele differences in HLA genes between 16th-century plague victims in Ellwangen, Germany and the modern local population, which could have been driven by selection (Immel et al., 2021); however, as with our study, they did not detect significant differences in HLA haplotype frequencies or diversity. While the signal of heterozygosity change has been recently used to map an ancient selective sweep in the HLA region (Souilmi et al., 2022) the fact that we were unable to detect significant changes in HLA heterozygosity in our sample may be a reflection of the short effect time of the pandemic.

Similarly, we did not find an excess of variants in genes related to health and immunity to be highly differentiated between the before and after the Black Death cohorts. The most highly differentiated variant in the *RIPK2* gene shows an increase in frequency after the Black Death of an allele protective against leprosy. Notably, this gene has been suggested to also have a role in the recognition of *Yersinia pestis* by the innate immune response (Ferwerda et al., 2009). Some scholars have suggested that the Second Plague Pandemic contributed to the decline of leprosy starting from the mid-13th century, as many people vulnerable to leprosy fell victim to the plague (Clay, 1909; Richards, 1977). On the other hand, we could not replicate in our Cambridge cohort the findings by Klunk et al. (Klunk et al., 2022) of significantly higher differentiation of immune genes, and, more specifically, differentiation of the four SNPs identified in their London and Danish cohorts (Table S4). The signal of enrichment disappears or even reverses direction when we use an extended set of neutral regions as our reference. The high correlation in allele frequency of common variants (MAF > 0.1 in Klunk et al. samples) between our Cambridge cohort and the London/Danish cohort, and the fact that we got qualitatively similar results when using the same variant lists suggest that differences in genotyping, either with or without imputation, are not likely the key factors; our results show that the subset of 250 regions used by Klunk et al. are likely to contain too few variants to be robustly representative of variation in the full list of 37,574 neutral regions defined by Gronau et al. 2011. Even though not supporting the results of Klunk et al. study, our results are in line with other research showing relatively low or moderate differentiation of immunity genes by Fst (Bronson et al., 2013; Maróstica et al., 2022) Low differentiation is consistent with the expectations of the theory of balancing selection (Brandt et al., 2018) as well as poison/antidote model (Enard & Petrov, 2018) whereby infectious disease transmission between communities often co-occurs with mobility and gene flow that locally increase the effective population size of genes involved in host defense.

Our limited sample size sets restrictions to identification of selection, yet, our sample size is comparable to the one used by Klunk et al. and even though low sequencing coverage could prevent us from identifying individual loci under selection the polygenic approach taken by Klunk et al., which we have followed, should in principle be able to detect selection at many variants contributing to a trait. Although our results revealed no enrichment of highly differentiated variants in immunity genes between the before and after the Black Death cohorts, these results do not mean that plague had no selective impact on genetic variation in Cambridge. The immune response to *Y. pestis* might involve multiple pathways which are yet to be fully understood and by our ‘blind’ approach of including all possible immune variants the signal of selection can be buried into background noise.

In sum, we have shown that even at very low coverage, whole genome sequencing of historical genomes when combined with rich historical context and other archaeological evidence can help to reconstruct many aspects of medieval life: the occasional incomers in a rural community; a young child from a benefactor family who was buried in the Friary; the Hospital full of unrelated inmates who might have traveled from afar; the social groups that had little difference in genetic background, but enjoyed different quality of life suggested by the stature they achieved; a pandemic whose genetic consequence in the Cambridge population remained evasive, despite the devastating effect on individual lives and medieval society. The range will surely expand with advancement in aDNA sequencing technology as well as framework to combine various sources of evidence.

## Acknowledgements

We are grateful to all our colleagues on the ‘After the Plague’ project, and to the Cambridge Archaeological Unit, Cambridgeshire County Council Historic Environment Team, and the Duckworth Laboratory for access to their collections and sampling permissions.

This research has been conducted using the UK Biobank Resource under Application Number 54698. Data analyses were carried out with the facilities of the High-Performance Computing Center of the University of Tartu.

## Authors contributions

R.H., C.L.S, J.E.R and T.K. conceived and designed the study; R.H., C.L.S., E.D’A., S.A.I., C.C., S.A.B., A.W.W., M.Q.A.A., S.J.G., A.S., H.N., X.J.G., A.K.R., O.B. performed data collection and analysis; R.H., C.L.S., E.D’A. and T.K. prepared the paper with critical input from all authors.

## Data and material availability

The ancient genomic data generated during this study are available at https://www.ebi.ac.uk/ena/browser/view/PRJEB59976 (accession code ENA: PRJEB59976) and the data depository of the EBC (https://evolbio.ut.ee/).

