## Supplementary material for "Medieval social landscape through the genetic history of Cambridgeshire before and after the Black Death": Materials and Methods and Supplementary Figures

### Supplementary Materials

**This PDF file includes:**

Materials and Methods

Figures S1 – S13

**Other Supplementary Materials for this manuscript include the following:**

Tables S1 – S13

### Materials and Methods

#### Sample information and ethical statement

All skeletal elements were sampled with permissions from the representative bodies/host institutions. Samples were taken and processed to maximize research value and minimize destructive sampling. Teeth were sampled from skeletons using gloves. Molars were preferred due to having more roots and larger mass, but premolars were also sampled.

#### Archaeological sites and material

Here we briefly describe the archaeological sites included in this study. More detailed information can be found at (Cessford et al., 2021).

*Cherry Hinton*

The settlement of Church End Cherry Hinton (Cherry Hinton) is located around six kilometers southeast of Cambridge. In the late 9th to the mid-10th century, a large *thengly* (aristocratic) or proto-manorial center was established (Cessford & Dickens, 2005b; Cessford & Slater, 2014). The associated timber chapel and graveyard were excavated in 1999 by the Hertfordshire Archaeological Trust (subsequently Archaeological Solutions and now Wardell Armstrong) (Lally, 2008).

Only part of the cemetery was investigated, including over 670 graves and the remains of *c.* 980 individuals. The cemetery population was estimated to be *c.* 1000–2000. The cemetery probably served an entire thriving Late Saxon to Norman rural settlement, broadly representative of a mixed rural peasant tenant hierarchy engaged primarily but not exclusively in agricultural labor. Based on radiocarbon dating plus a range of other evidence the cemetery dates to *c.* 940/990–1120/70. As the population from which those buried at Church End were drawn probably increased over time it is likely that the majority, possibly around two thirds, died after the Norman Conquest. As this settlement was located firmly within the rural hinterland of Cambridge this population provides an important comparator to the town, albeit one that is rather earlier than the main populations that have been studied from there.

*All Saints*

All Saints by the Castle (All Saints) is a medieval parish church and associated cemetery in Cambridge. Although not documented until 1217, the church and cemetery are likely to have been founded *c.* 940–1100 and the parish was amalgamated with St Giles in 1365/6 (HMC, 1877) (299, fol. 1026).

The cemetery of All Saints was first identified in 1972 in evaluation trenches by John Alexander, then partially investigated by Paul Craddock and Vince Gregory in 1973 (Craddock & Gregory, 1975), with further excavations in 1988 and 1994 (Cessford & Dickens, 2005a). In total just over 210 skeletons have been excavated from the cemetery, out of an estimated original overall total of *c.* 2500–3500. The burials at the cemetery of All Saints probably constitute a broadly representative sample of the parishioners' *c.* 940/1100–1365/6, albeit slightly distorted by several factors. Most burials, probably over 80%, are likely to date to the period after the Norman Conquest. Although the population of All Saints was socially and economically mixed, they are probably somewhat poorer than the average for Cambridge and rather more involved in agricultural activities. Radiocarbon dating is compatible with the burials dating to c. 940/1100–1365/6. The burials at All Saints begin and end earlier than the other main Cambridge urban skeletal assemblages, from the Hospital of St John and the Augustinian Friary.

*Hospital of St John*

The Hospital of St John the Evangelist in Cambridge (Hospital or Hospital of St John) was established by local townspeople *c.* 1190–1200 to care for, principally in a social and spiritual sense rather than medically, the poor, infirm and sick. It did this until it was dissolved to create St John's College in 1511, although it was allegedly already in decline during the 15th century. Excavations in 2010–2011, conducted by the Cambridge Archaeological Unit in the detached cemetery of the hospital revealed almost 400 complete or partial skeletons, from a likely original population of *c.* 1000–1500 CE burials (Cessford, 2015).

The results of the intensive radiocarbon dating program and other evidence are compatible with burial starting in the early 13th century and indicate that it continued until at least the mid/late 15th century.

The burials at the detached cemetery of the Hospital of St John represent an amalgam of various groups associated with the institution. Most of them were probably charitable inmates of the Hospital.

*Augustinian Friary*

The Augustinian Friary (Friary) was established in Cambridge between 1279/80 and 1289, when it was first mentioned in a royal pittance. The Friary grew rapidly and thrived, becoming a *studium generale* or national study house with internal connections in 1318 and having 70 friars present in 1328. It continued until the Dissolution 1538, as one of the most important Augustinian friaries in England and one of the largest institutions in Cambridge.

Human remains have been recovered from three locations at the Friary: an early cemetery and the later chapter house and cloister. Burials included in this study were excavated by the Cambridge Archaeological Unit in 2016–2017 (Cessford, 2017, 2020; Cessford & Neil, 2022). Some of the burials were accompanied by single buckles located near the pelvis, indicating that the bodies were buried in a clothed state, with surviving evidence for associated leather girdles and some evidence for textiles. Some of the skeletons definitely lacked buckles and these were probably buried in shrouds. It appears that members of the Augustinian order received clothed burial, while shrouded burials are of lay individuals (Cessford & Neil, 2022). The lay individuals would include patrons and benefactors, as well as lay servants of the Friary and corrodians. The 72 burials recovered from the Friary represent only a portion of the estimated 200–700 individuals likely to have been interred there between 1290 and 1538.

*Bene't Street*

St Bene't's (a contraction of Benedict's) parochial church in Cambridge was established in *c.* 1000–1050 CE and remains in use, with burial continuing until the 1850s. A strip of land along the western side of the churchyard was transferred to Corpus Christi College between 1352 and 1377, to form an entrance route between Bene't Street and the College. Part of this strip was excavated by the Cambridge Archaeological Unit in 2006, revealing highly truncated individual burials and part of one mass burial (Cessford & Fallon, 2006). Among four skeletons tested from the mass burial, two yielded positive and one tentative identifications of *Y. pestis* (Cessford et al., 2021). It is probable that this mass burial relates to the Black Death as it is likely to predate the construction of the College buildings starting in 1352.

*Baptist Chapel*

The Providence Calvinistic Baptist Chapel (Baptist Chapel) in Cambridge was in use for just four years 1833–1837 CE. Part of the associated cemetery was excavated in 2012, by Oxford Archaeology East (Rees, 2014). There originally may have been a maximum of 20 graves in the investigated area, with at least 16 graves in five rows 11 of which were recorded. The graves were aligned north-northwest to south-southeast and were mainly earth cut, but two were brick-lined and there is evidence for coffins and shrouds.

*Holy Trinity*

Holy Trinity Church in Cambridge (Holy Trinity) is first mentioned in 1174, when it is said to have been burnt down, and was probably established *c.* 1050–1150 CE. Excavations by the Cambridge Archaeological Unit in 2016–17 revealed seventeen articulated burials (Newman, 2018). Although these were reburied, they were subject to thorough osteological analysis and sampling. The skeletons included in this study were placed in coffins in the vaults constructed after a vestry building of 1833/4 and prior to the end of burial in the cemetery in 1855.

*Midsummer Common*

From 1574 until the last outbreak in Cambridge in 1665/6, some individuals infected with the plague were isolated and moved to pest houses, located some distance from the town in its surrounding fields. Although primarily about quarantine and isolation of the living, parish registers record that from at least 1603 onwards some individuals who died at pest houses were buried there rather than returned to their parish cemetery. The location of the pest houses changed over time, with at least four locations known. One of these sites was Midsummer Common, where pest houses are mentioned in 1593 and 1630. Two skulls discovered in the late 19th or early 20th century at Midsummer Common were presented to the Duckworth Collection by the town clerk, John Edleston Ledsam Whitehead (1853–1923, town clerk 1887–1923). A tooth from one of the skulls was dated to 1450–1630 cal ad (Cessford et al., 2021).

*Clopton*

Clopton was a village in west Cambridgeshire, about 19 kilometers southwest of Cambridge. The village was established by the 10th–11th centuries and appears to have thrived until forcible enclosure for sheep grazing in *c.* 1480–1520 CE. The church is documented as having gone out of use in 1561. There is good evidence that the village fell primarily within the hinterland of Cambridge, although its location meant that it also had links to urban centers in Hertfordshire. The church of St Mary had been established at Clopton by the late twelfth century, but is probably significantly older. A new church was dedicated in 1352 and this continued to stand after the village was deserted.

John Alexander excavated *c.* 70 skeletons from the church and cemetery between 1960 and 1964, out of over 120 burials with records (Alexander, 1968). The excavations focussed primarily on the cemetery to the south of the church, which would have been where the bulk of the parishioners were buried. The limited excavations in and around the church means that the individuals who were buried there, which would have included clerics such as John Thorney as well as wealthy parishioners, are largely absent from the studied human remains. Radiocarbon dating and other evidence is consistent with a date of *c.* 1200–1561 CE for the burials.

Hemingford Grey

Excavations at Meadow Lane, north of Hemingford Grey, 19km northwest of Cambridge by Oxford Archaeology in 2006 revealed sixteen burials in a late 17th–early 18th-century nonconformist cemetery that has been linked to the Society of Friends or Quakers (Sims, 2007). Textual evidence for the cemetery comes from the vicar at Hemingford Grey Parish noting in the Parish register burials that took place away from the Parish cemetery at a site referred to as Wobourn or Oubourne. This lists 17 burials between 1681 and 1721, although it is unclear if the list is complete and earlier and later burials are possible. The individuals buried came from Hemingford Grey and the adjacent Parishes of Fenstanton, St Ives and Hemingford Abbot and there are several examples of multiple members of a single family.

Dating

The radiocarbon determinations were undertaken at the Scottish Universities Environmental Research Centre (SUERC) radiocarbon laboratory and followed their standard procedures (Dunbar et al., 2016). Analysis was undertaken using OxCal v.4.3 (Bronk Ramsey, 2009; Bronk Ramsey & Lee, 2013) using the IntCal13 calibration curve (Reimer et al., 2013). A small number of additional radiocarbon determinations (marked by * in Table S1) were undertaken at a late stage, to date specific individuals that produced evidence for specific pathogen aDNA. These were undertaken at the 14Chrono Centre, Queen's University Belfast. These additional determinations were calibrated using IntCal 2020 (Reimer et al., 2020).

At the Hospital of St John, Augustinian Friary and All Saints by the Castle, detailed Bayesian modeling of these results combined with expert archaeological judgment has helped to inform whether these individuals (along with others in the stratigraphic sequence) died before or after the Black Death pandemic of 1348–9. More details can be found in (Cessford et al., forthcoming).

Sampling, ancient DNA extraction and library preparation

Inside a class IIB hood in the dedicated aDNA facility of the University of Cambridge Department of Archaeology or University of Tartu Institute of Genomics, root portions of teeth were removed with a sterile drill wheel. Petrous was sampled with a 10mm core drill sterilized with bleach followed by distilled water and then ethanol rinse. Root and petrous portions were briefly brushed to remove surface dirt, any varnish or lacquer, and microbial film with full strength household bleach (6% w/v NaOCl) using a disposable toothbrush that was soaked in 6% (w/v) bleach prior to use. They were then soaked in 6% (w/v) bleach for 5 minutes. Samples were rinsed twice with 18.2 MΩcm H2O and soaked in 70% (v/v) Ethanol for 2 minutes, transferred to a clean paper towel on a rack inside a class IIB hood with the UV light on and allowed to dry. They were weighed and transferred to PCR-clean 5 ml or 15 ml conical tubes (Eppendorf) for chemical extraction.

Inside a class IIB hood, per 100 mg of each sample, 2 ml of 0.5M EDTA Buffer pH 8.0 (Fluka) and 50 µl of Proteinase K 10 mg/ml (Roche) was added. Tubes were rocked in an incubator for 72 hours at room temperature. Extracts were concentrated to 250 µl using Amplicon Ultra-15 concentrators with a 30 kDa filter (Millipore). Samples were purified according to manufacturer's instructions using buffers from the Minelute™ PCR Purification Kit (Qiagen) with the following changes: 1) the use of High-Volume spin columns (Roche); 2) 10X PB buffer instead of 5X; and 3) samples incubated with EB buffer (Qiagen) at 37C for 10 minutes prior to elution. The columns were transferred to clean, labeled, 1.5ml Eppendorf tubes. One hundred microlitres EB buffer is added to the membrane and centrifuged at 13,000 rpm for two minutes after the 10-minute incubation and stored at -20 C. Only one extraction was performed per sample for screening and 30µl used for libraries.

Library preparation was conducted using a protocol modified from the manufacturer's instructions included in the NEBNext® Library Preparation Kit for 454 (E6070S, New England Biolabs, Ipswich, MA) as detailed in (Meyer & Kircher, 2010). DNA was not fragmented and reactions were scaled to half volume, adaptors were made as described in (Meyer & Kircher, 2010) and used in a final concentration of 2.5uM each. DNA was purified on MinElute columns (Qiagen). Libraries were amplified using the following PCR set up: 50µl DNA library, 1X PCR buffer, 2.5mM MgCl2, 1 mg/ml BSA, 0.2µM inPE1.0, 0.2mM dNTP each, 0.1U/µl HGS Taq Diamond and 0.2µM indexing primer. Cycling conditions were: 5' at 94C, followed by 18 cycles of 30 seconds each at 94C, 60C, and 68C, with a final extension of 7 minutes at 72C. Amplified products were purified using MinElute columns and eluted in 35 µl EB (Qiagen). Three verification steps were implemented to make sure library preparation was successful and to measure the concentration of dsDNA/sequencing libraries – fluorometric quantitation (Qubit, Thermo Fisher Scientific), parallel capillary electrophoresis (Fragment Analyser, Advanced Analytical) and qPCR.

DNA sequencing

DNA was sequenced using the Illumina NextSeq500/550 High-Output single-end 75 cycle kit at the University of Cambridge Department of Biochemistry DNA Sequencing Facility. As a norm, 20 samples were sequenced together on one flow cell; additional data was generated for samples over 5% endogenous human content to increase coverage.

Mapping

Before mapping, the sequences of the adapters, indexes, and poly-G tails were removed from read ends and reads shorter than 30 bp were removed using cutadapt-1.11 (Martin, 2011).

The sequences were aligned to the reference sequence GRCh37 (hg19) using Burrows-Wheeler Aligner (BWA 0.7.12) (Li & Durbin, 2009) and the command *aln* with re-seeding disabled.

After alignment, the sequences were converted to BAM format and only sequences that mapped to the human genome were kept with samtools 1.3 (Li et al., 2009). Data from different flow cell lanes were merged and duplicates were removed using picard 2.12 (http://broadinstitute.github.io/picard/index.html).

Sequencing coverage was estimated using Qualimap 2.2.1 (Okonechnikov et al., 2015) after applying a custom accessibility mask that includes the 1000 genomes accessibility mask and a composite mappability track (Derrien et al., 2012).

aDNA authentication

As a result of degradation over time, aDNA can be distinguished from modern DNA by certain characteristics: short fragments and a high frequency of C > T substitutions at the 5' ends of sequences due to cytosine deamination. The program mapDamage2.0 (Jónsson et al., 2013) was used to estimate the frequency of 5' C > T transitions. Rates of contamination were estimated from mtDNA using ContamMix (Fu et al., 2013) and from X chromosome using two methods implemented in ANGSD (Korneliussen et al., 2014).

Samtools 1.3 (Li et al., 2009) option *stats* was used to determine the number of final reads, average read length, average coverage etc. The average endogenous DNA content (proportion of reads mapping to the human genome) was 13.22% (0.00% - 81.27%).

Calculating genetic sex estimation

Genetic sex was calculated using the methods and script described in (Skoglund et al., 2013), estimating the fraction of reads mapping to Y chromosome out of all reads mapping to either X or Y chromosome. Genetic sex was calculated for samples with a coverage >0.01x and only reads with a mapping quality >30 were counted for the autosomal, X, and Y chromosome.

Determining mtDNA haplogroups

Mitochondrial DNA haplogroups were determined using Haplogrep2 (Weissensteiner et al., 2016). Subsequently, the identical results between the individuals were checked visually by aligning mapped reads to the reference sequence using samtools-1.3 (Li et al., 2009) command *tview* and confirming the haplogroup assignment in PhyloTree (accessed at: www.phylotree.org). Additionally, private mutations were noted for further kinship analysis.

Y chromosome variant calling and haplotyping

A total of 256,463 binary Y chromosome SNPs that have been detected as polymorphic in previous high coverage whole Y chromosome sequencing studies (Hallast et al., 2015; Karmin et al., 2015; Poznik et al., 2016) and by YFull (https://www.yfull.com/snp-list/) were called in male samples with more than 0.001× autosomal coverage using ANGSD-0.916 (Korneliussen et al., 2014) '-doHaploCall' option. Basal haplogroup affiliations (Table S1) could be determined for 128 samples by assessing the proportion of derived allele calls (pD) in a set of primary (A, B, C...T) haplogroup defining internal branches, as defined in (Karmin et al., 2015), using 4081 informative sites. Haplogroup assignments were confirmed and specified using pathPhynder (Martiniano et al., 2022). Further detailed sub-haplogroup assignments within the phylogeny of the primary haplogroup were determined on the basis of mapping the derived allele calls to the internal branches of the YFull tee (https://www.yfull.com/tree/), requiring support of at least two variants for the terminal branch assignment.

Pseudohaploid genotype calling

Autosomal variants were called with the ANGSD 0.917 (Korneliussen et al., 2014) command `--doHaploCall`, which randomly selects one base at each specified position in the genome. The pseudohaploid genotypes were used in genetic kinship analysis with READ and for PC analyses of individual sites with smartPCA (Figures S1-9).

Imputation

Following (Hui et al., 2020), genotype likelihoods were first updated with Beagle 4.1 (Browning and Browning 2016) from genotype likelihoods produced by ANGSD 0.917 (`-doMajorMinor 3 -GL 1 -doPost 1 -doVcf 1 -sites …`) (Korneliussen et al., 2014) in Beagle -gl mode, followed by imputation in Beagle -gt mode with Beagle 5 (Browning et al., 2018) from sites where the genotype probability (GP) of the most likely genotype reaches 0.99. To balance between imputation time and accuracy, we used 503 Europeans genomes in 1000 Genomes Project Phase 3 (The 1000 Genomes Project

Consortium, 2015) as the reference panel in Beagle -gl step, and 27,165 genomes (except for chromosome 1, where the sample size is reduced to 22,691 due to a processing issue in the release) from the Haplotype Reference Consortium (HRC) (McCarthy et al., 2016) in the Beagle -gt step. After filtering out rare variants with MAC <5, we retained a total of ~37 million variants genome-wide. Because Beagle treats "./." in the VCF input as sporadically missing and imputes them during haplotype phasing, which damages the accuracy when such missing genotypes are common, we imputed each genome individually so that missing genotypes were not included in the VCF input to Beagle 5. The output single-sample VCFs were then merged for downstream analysis. Apart from newly generated genomes in this study, we also imputed published genomes from (Margaryan et al., 2020; Martiniano et al., 2016; Schiffels et al., 2016).

We down-sampled a 10.83x genome that has been previously reported in (Guellil et al., 2022), PSN31, to 0.05x-1x to evaluate our imputation pipeline (Table S13). The accuracy was estimated by comparing the imputed genotypes to genotypes called using GATK HaplotypeCaller (McKenna et al., 2010) (with the flags `--min-pruning 1 --min-dangling-branch-length 1`). For some analysis, we filtered the imputed genotypes further by GP and/or minor allele frequencies at the sites as described in the respective sections.

Principal component analysis

We used FlashPCA2 (Abraham et al., 2017) for principal component analysis (PCA) of imputed genomes (without projection) together with modern reference genomes from selected groups in UK Biobank after excluding variants in linkage disequilibrium with the PLINK --indep-pairwise 1000 50 0.5 option and exclusion of the likely non-neutral regions exclusion_regions_hg19.txt (Figure 1C and Figure S1). For comparison we also performed PCA using modern reference genomes only, before projecting pseudo-haploid genomes (coverage > 0.05X) onto the PC space following (https://github.com/chrchang/eigensoft/blob/master/POPGEN/lsqproject.pdf) (Figures S2-11). The error bars of the projected genomes represent one standard deviation obtained from 20-fold block jackknife.

Detecting IBD and long shared allele interval (LSAI) segments

IBD/LSAI segments (Kivisild et al., 2021) and kinship coefficients were estimated from merged plink files of 61 imputed ancient genomes, 503 Europeans from the 1000 Genome Project and UK Biobank data with IBIS version 1.20.9 (Seidman et al., 2020) using minimum shared segment length (-min_L) thresholds 5 cM and/or 7 cM together with -maxDist 0.1 and -mt 300 parameters. In total, 269,319 binary SNPs with MAF > 0.05 were used.

Although IBIS has the highest IBD inference accuracy for > 7 cM segments, which we use in kinship analyses, we use > 5 cM threshold in our diachronic inferences of population affinities because our focus is on relationships at generational distances > 15 at which longer IBD sharing expectations become relatively low, particularly in combination with the loss of sensitivity to detect long IBD segments from imputed ancient DNA sequences. Because true IBD segments of this length are not expected to be common at these generational distances, we need to consider the detected segments as "long shared allele intervals" (LSAIs) rather than IBD segments sensu stricto. Because they are inferred from unphased data after removal of rare variants (which cannot be imputed with sufficient accuracy), the LSAIs are likely to include undetected recombination points and smaller IBD segments residing on different haplotypes.

The probability of individual connectedness (PiC) score (Kivisild et al., 2021) for individual x in group Z was estimated as the proportion of individuals from group Z with whom individual x shared IBD above the given threshold. In practice, we estimated the count of connected individuals from group Z from sorted IBIS .coef output files by using the linux "join" function to add group codes to individual identifiers and by using the "crosstab" function of datamash[59] to generate the table of counts, each of which we divided by the total number of individuals in group Z to obtain the individual connectedness proportions by groups (the PiC scores).

Unsupervised community extraction analyses on IBIS .coef files calculated from merged plink files including imputed ancient genomes with coverage > 0.2x and modern UK Biobank data were run with the Louvain algorithm (Blondel et al., 2008) implemented in the R library "igraph" (Csardi & Nepusz, 2006) with additional significance tests as described in (Kivisild et al., 2021) using scripts available at https://github.com/SABiagini/Louvain.

Genetic kinship analysis

Pseudohaploid genotypes at 5,494,912 positions with MAF > 0.05 in the UK10K subset of the HRC panel were called with ANGSD version 0.917. For the comparison with published studies, the merged PLINK file from the PCA analysis was used and select populations retained using `plink --keep` and converted to .tped. The ANGSD output files were converted to .tped format, which was used as an input for kinship analyses with READ (Kuhn et al., 2018). Given highly homogenous ancestry in all study sites, samples from all sites were analyzed together for Figure 2 as well as separately (results not shown) to test for potential biases in normalized P0 calculation. In addition to 1st and 2nd degree relationships we also estimated P0 cutoffs (15/16 = 0.9375 as per (Kuhn et al., 2018)) for the detection of 3rd degree relatives while acknowledging that due the lack of relevant empirical data the false positive and negative error rates for this class of relationship remain unknown.

Runs of homozygosity

We used hapROH (Ringbauer et al., 2021) to detect runs of homozygosity (ROH) in ancient genomes. Using information from a reference panel, hapROH has been shown to work for genomes with more than 400K of the 1240K SNPs panel covered at an error rate lower than 3% in pseudo-haploid genotypes (Ringbauer et al., 2021). We note that the requirement is broadly in line with the imputation accuracy we get from coverages as low as 0.1x, where ~60% of common variants (MAF≥0.05) in the HRC panel are recovered with an overall accuracy around 97% for diploid genotypes (Table S13). Among common variants in the HRC panel, 853,159 overlap with the 1240K SNPs panel.

1000 Genomes Project data were used to construct the reference haplotypes. We kept the standard parameters in hapROH, which had been optimized for 1240K aDNA genotype data:

```
e_model='haploid', post_model='Standard', random_allele=True,
roh_in=1, roh_out=20, roh_jump=300, e_rate=0.01, e_rate_ref=0.0,
cutoff_post=0.999, max_gap=0, roh_min_l=0.01
```

Heterozygosity

Genome-wide heterozygosity and heterozygosity in the HLA region (--chr 6 --from-kb 28477 --to-kb 33448) were estimated with the --het function in PLINK 1.9 (Chang et al., 2015) for imputed genomes with >0.1x coverage and for 5,448,740 sites that had MAF > 0.05 in the

HRC reference panel. Sites where the highest genotype probability is less than 0.99 were set to missing.

Nucleotide diversity

Nucleotide diversity in the HLA region and throughout the autosomes was estimated using vcftools --window-pi (Danecek et al., 2011), which by default outputs result in 10kb windows, in imputed genomes with > 0.05x coverage. Sites where the highest genotype probability is less than 0.99 were set to missing. The results are similar whether we filtered the variants by MAF > 0.05 in the HRC reference panel or not (Figure S13).

Stature estimation

Long bone measurements (clavicle, humerus, radius, ulna, femur, tibia and fibula) were taken as outlined in Buikstra and Ubelaker (Buikstra & Ubelaker, 1994). Measurements were taken using an osteometric board and sliding callipers. Circumferences were taken using a measuring tape. All measurements were taken to 1pd. When left and right bones were measurable, a mean of the two measurements was calculated. Measurements were not taken for the purpose of stature reconstruction for individuals with gross pathologies (e.g. vitamin D deficiency), or from bones with mal-aligned trauma which altered the length of the bone. Only individuals who were developmentally complete (long bone epiphyses fused) were measured for stature estimation. Published measurements were taken for Clopton and the material excavated from the Friary in the 1900s as it was no longer possible to access the post crania of these remains. The appendices in (Inskip, forthcoming) outline all measurements taken.

Stature was calculated from long bone measurements following Trotter (1970) as included in (White & Folkens, 2005). These formulae were selected as many of the comparative collections analyzed in the mid 20th century, for which post crania are no longer available, used these equations. Due to the level of truncation at many of the sites assessed by the ATP project, it was decided to use single bone estimates rather than those that required two bones (e.g. femur and tibia) to maximize sample size. Stature estimates were not derived from the fibulae due to the low number of complete bones. Stature estimated from various long bones can also be found in the appendices of (Inskip, forthcoming).

Polygenic risk score for height

Eight polygenic risk score (PRS) models for human height trained on UK Biobank data were applied on the imputed genomes: PRS, LDpred, MCP, Lasso, PRS+MTAG, LDpred+MTAG, MCP+CTPR and Lasso+CTPR (Chung et al., 2019). Some of the models (MTAG and CTPR) also incorporated information about BMI to aid modeling height. The authors have shown that these models can reach prediction $R^2$ between 0.22 and 0.43 in modern validation datasets. Coefficients of the models were downloaded from http://lianglab.rc.fas.harvard.edu/CTPR/.

Each model was applied on two versions of the imputed genotypes: dosage data without further filtering, and hardcall genotypes after filtering by GP (max(GP) ≥ 0.99). We calculated the polygenic risk scores from each linear model in PLINK 1.9 (using `--score` flag) (Chang et al., 2015).

Table S9 shows the correlation coefficients ($R^2$) between the PRS calculated using different models and stature estimated from long bones in our samples from the later medieval period. We note that when low coverage genomes (< 0.1x) are included, where the imputed genotypes carry more uncertainty, PRS calculated from the dosage data show higher correlation to stature estimates. The correlation also increases when we raise the coverage threshold, and when we only include stature estimated from the femur. We chose to include 88 genomes above 0.05x with stature estimated from the femurs as a trade-off between the sample size and the predictive power of the PRS to fit the regression models.

Regression models for stature

We fitted a range of Bayesian linear regression models using stature estimation as the outcome variable. The models take the general form

$$\mu = \beta \cdot \mathbf{X} + \alpha$$

$$\mathbf{y} \sim \mathrm{N}(\mu, \sigma^2)$$

Where **y** denotes femur based stature estimation; **X** denotes the matrix of numeric predictor variables, if any; $\beta$ is the vector of coefficients; $\alpha$ is the vector of intercept. There are multiple $\alpha$ terms when categorical predictor variables (e.g. sex or social group) are included in the model. Both **X** and **y** were scaled to have mean 0 and standard deviation 1. The prior distribution of the parameters were set to

$$\alpha \sim \mathrm{N}(0, 1)$$

$$\beta \sim \mathrm{N}(0, 0.5)$$

$$\sigma \sim \exp(1)$$

The “rstan” package (Stan Development Team, 2023) in R (R Core Team, 2022) was used to fit the regression models, and the “loo” package (Vehtari et al., 2022) was used for model comparison. The predictors we included in each model are: 1. Sex only; 2. Sex and PRS; 3. Sex and social group; 4. Sex, social group, and PRS. Since we would like to compare the fitted models, we only included 76 later medieval individuals supported by sufficient contextual information to belong to one of the four social groups: charity cases (a subset of burials from St John's Hospital); friars (a subset from Augustinian Friary); ordinary urban (burials from the parish cemetery of All Saints); ordinary rural (burials from the church at Cherry Hinton). Model 2 was separately fitted to all 88 genomes sequenced to > 0.05x and have stature estimates from the femur to produce Figure 4A.

The posterior distributions of each model we fitted were summarized in Table S10. All models were run with 4 chains of 50,000 iterations (including 25,000 burn-in iterations) each. The R-hat statistic was monitored to check the chains have reached convergence; we also checked that the effective sample size of all parameters is at least around 20,000. When the four models were compared based on the theoretical expected log pointwise predictive density, the models including PRS perform better than those without PRS (Table S10). The model without social group in the predictors (Model 2), appears to outperform the one including social group (Model 4) slightly, but the difference is not conclusive given the standard error of the difference.

To investigate the difference in stature between social groups when controlling for sex and PRS, we extracted the posterior samples from Model 4 and looked at the distribution of the difference between the intercepts associated with social groups (Figure 4B).

For sensitivity analysis, we changed to more diffused prior distributions for all parameters (N(0, 2) for $\alpha$; N(0, 1) for $\beta$; exp(0.5) for $\sigma$) and reran the four models. This led to more dispersion in the posterior distributions of $\alpha$ (the intercepts associated with indicator variables for sex and social groups), but our conclusions about model comparison and social group differences remained robust.

Phenotype prediction

Using the imputed genomes generated in this study (143 individuals with a coverage higher than 0.05x) and previously published (71 samples) (Margaryan et al., 2020; Martiniano et al., 2016; Schiffels et al., 2016), we extracted genotype calls for 39 out of the 41 HIrisPlex-S variants and 74 SNPs involved in diet and diseases, coding the allele information as the number of the effective allele (0, 1, 2) using plink (Tables S4-6). The diet and disease set of 74 SNPs was selected starting from lists of variants previously analyzed in ancient DNA studies (Allentoft et al., 2015; Günther et al., 2018; Olalde et al., 2014), prioritizing those with a role in the response to pathogenic infection. In addition to the 70 variants that we have previously used for phenotype prediction (Saag et al., 2021; Saupe et al., 2021), we also analyzed the 4 SNPs in immune loci recently detected to be highly differentiated before or after the Second Pandemic (Klunk et al., 2022) (Table S6). A table with the HIrisPlex-S SNP alleles per sample was uploaded on the HIrisPlex-S webtool (https://hirisplex.erasmusmc.nl/) to obtain probabilities values for each eye, hair and skin color category per sample. This output was then interpreted following the manual to obtain the final pigmentation prediction for each individual (Table S5). We then grouped the individuals by time period. The groups were compared performing an ANOVA statistical test, applying a Bonferroni's correction on each group of variants (carbohydrate metabolism, lipid metabolism, vitamin metabolism, response to pathogens, autoimmune diseases, other diseases, Pigmentation) (Table S4) to set the significance threshold. For only the significant SNPs, we also performed a *post-hoc* Tukey test to identify the significantly different group pairs. For our medieval samples, we also grouped them by relative time to the Second Pandemic (before or after) and performed a statistical t-test as reported in Table S4.

To test the enrichment for higher allele frequency differences at variant positions of immune genes we used Weir and Cockerham (Weir & Cockerham, 1984) Fst as implemented in PLINK (--fst case-control) on lists of variants derived from the Klunk et al. (Klunk et al., 2022) study, as well as an expanded list of 54,931 variants polymorphic in the HRC panel from the 37,574 putatively neutral regions defined by Gronau et al. 2011 that had MAF > 0.1 in 83 imputed genomes from Cambridge dating to before and after the Black Death. As an expanded set of immunity regions, we extracted 19,940 variants with MAF> 0.1 in the 83 Cambridge cohort from 189,173 exonic regions of the 4,723 innate immunity genes curated by InnateDB, https://www.innatedb.com/moleculeSearch.do.

### Supplementary Figures

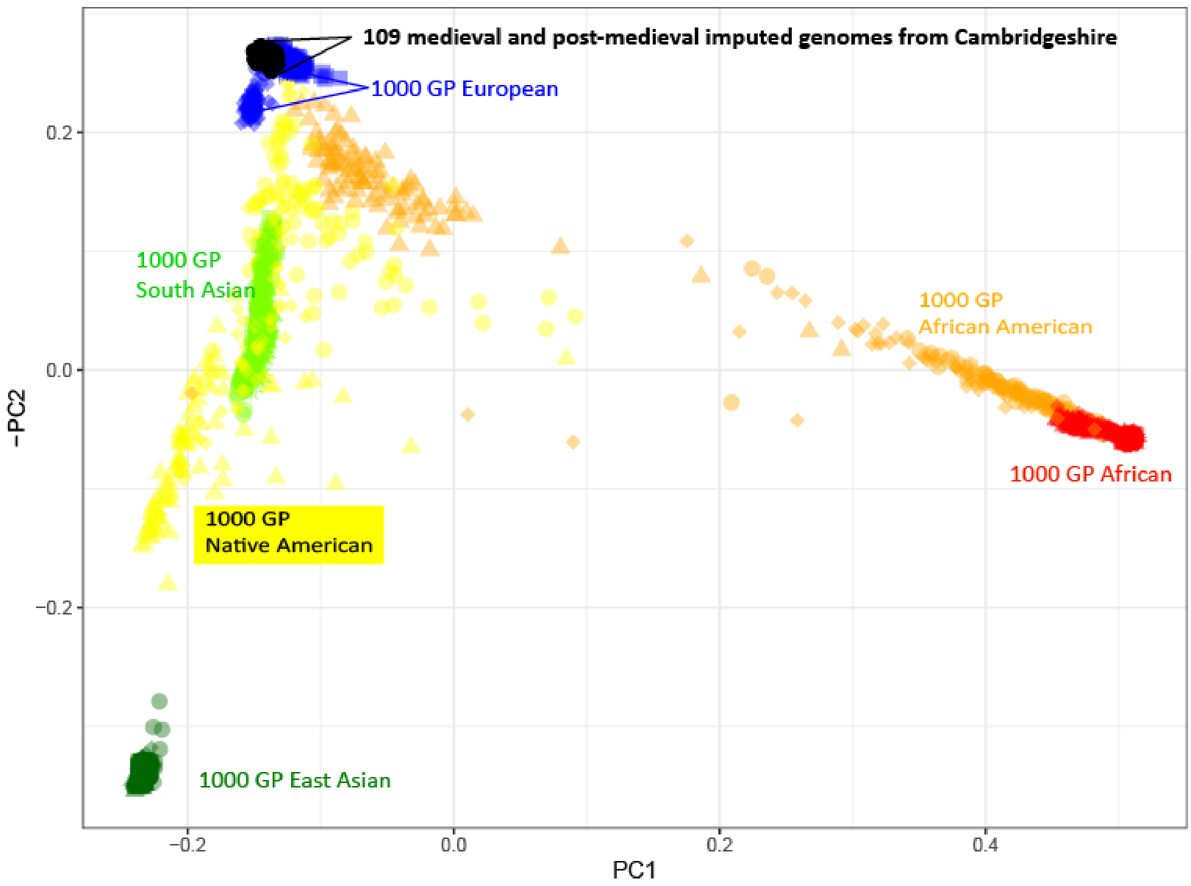


**Figure S1. PCA of 109 later medieval and post-medieval genomes (>0.1x) in context of 1000 Genome Project (1000 GP) data.**

**Figure S2-S11. PCA of West Europe individuals from UK Biobank, showing lsq-projection of historical genomes from each site.** Only genomes > 0.05x are included; error bars represent one standard deviation estimated from 20-fold block jackknife.

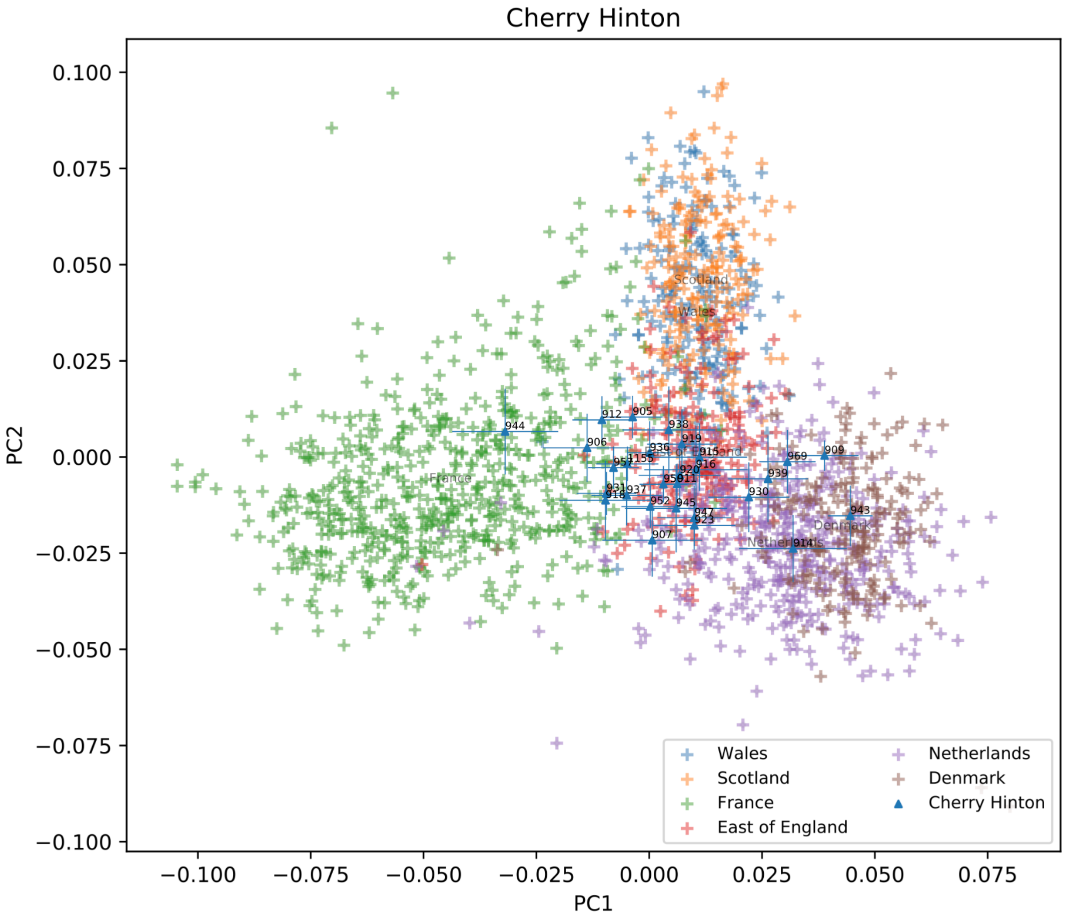


**Figure S2: PCA of West Europe individuals from UK Biobank, showing lsq-projection of historical genomes from Cherry Hinton.**

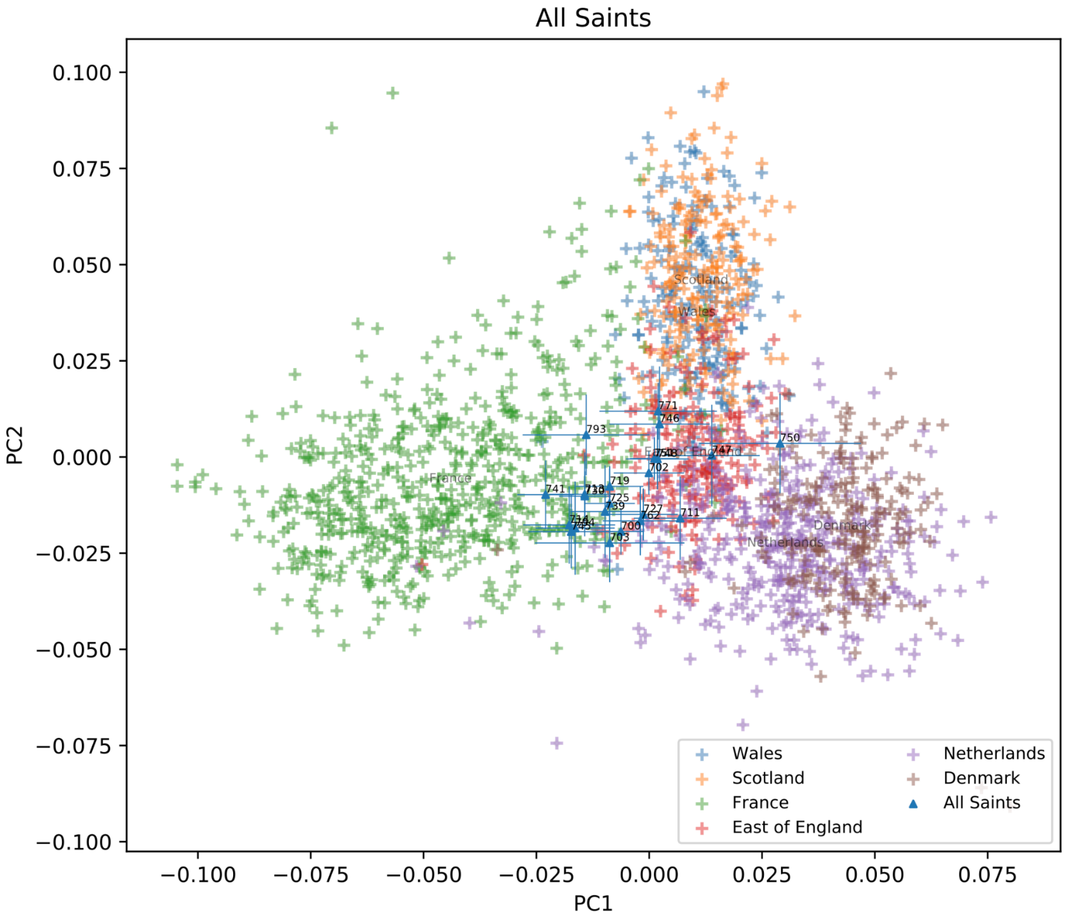


**Figure S3: PCA of West Europe individuals from UK Biobank, showing lsq-projection of historical genomes from All Saints by the Castle.**

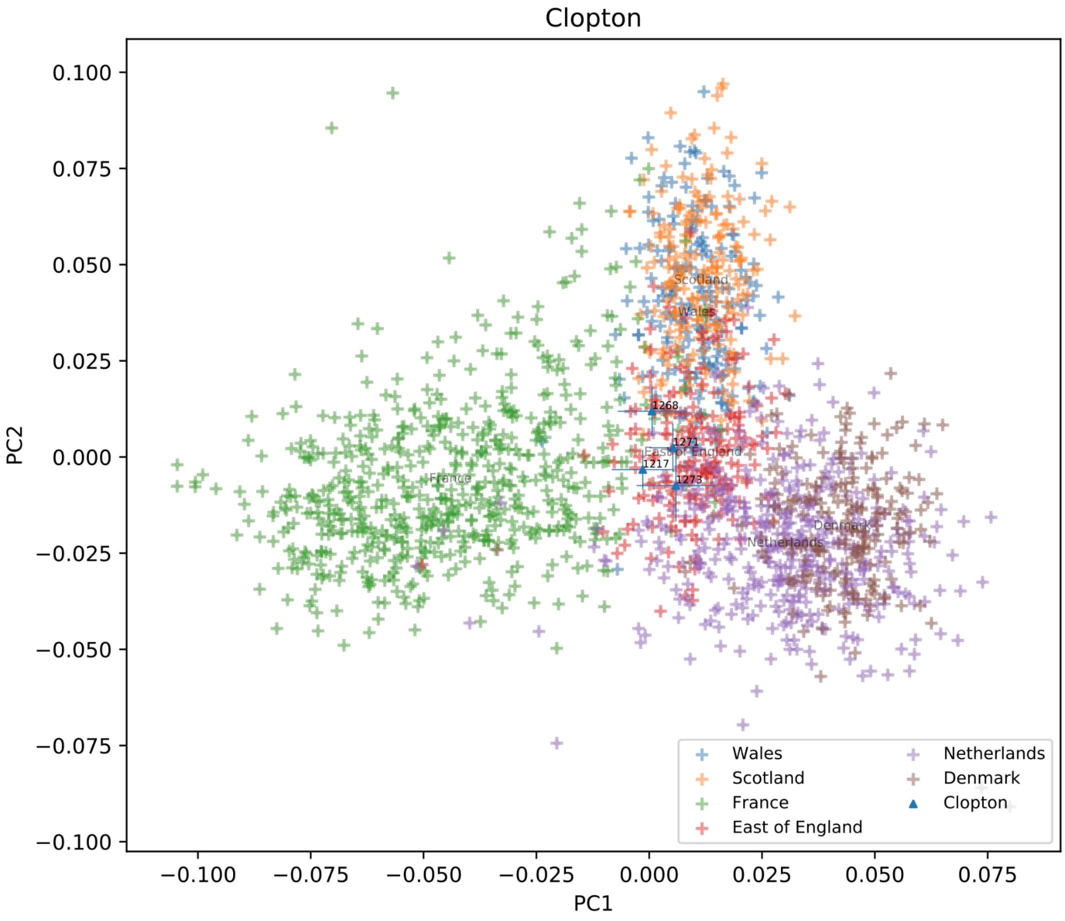


**Figure S4: PCA of West Europe individuals from UK Biobank, showing lsq-projection of historical genomes from Clopton.**

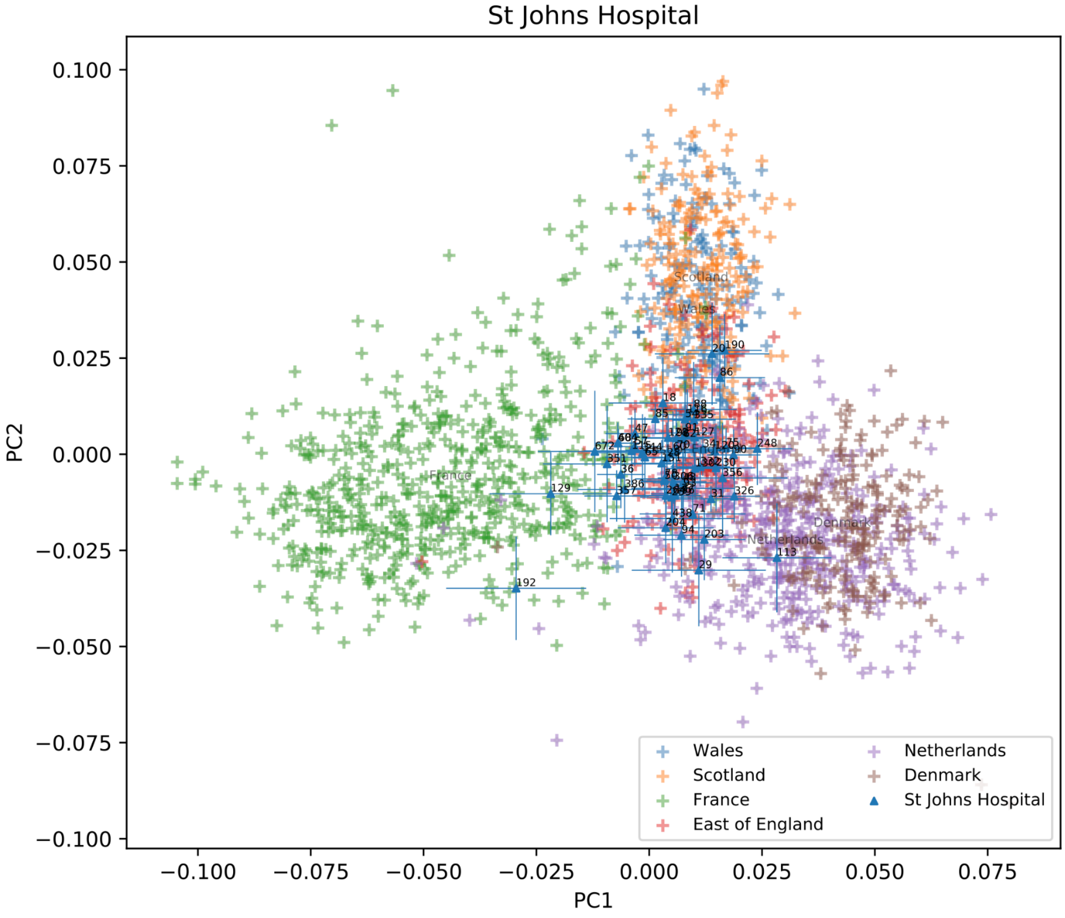


**Figure S5: PCA of West Europe individuals from UK Biobank, showing lsq-projection of historical genomes from the Hospital of St John.**

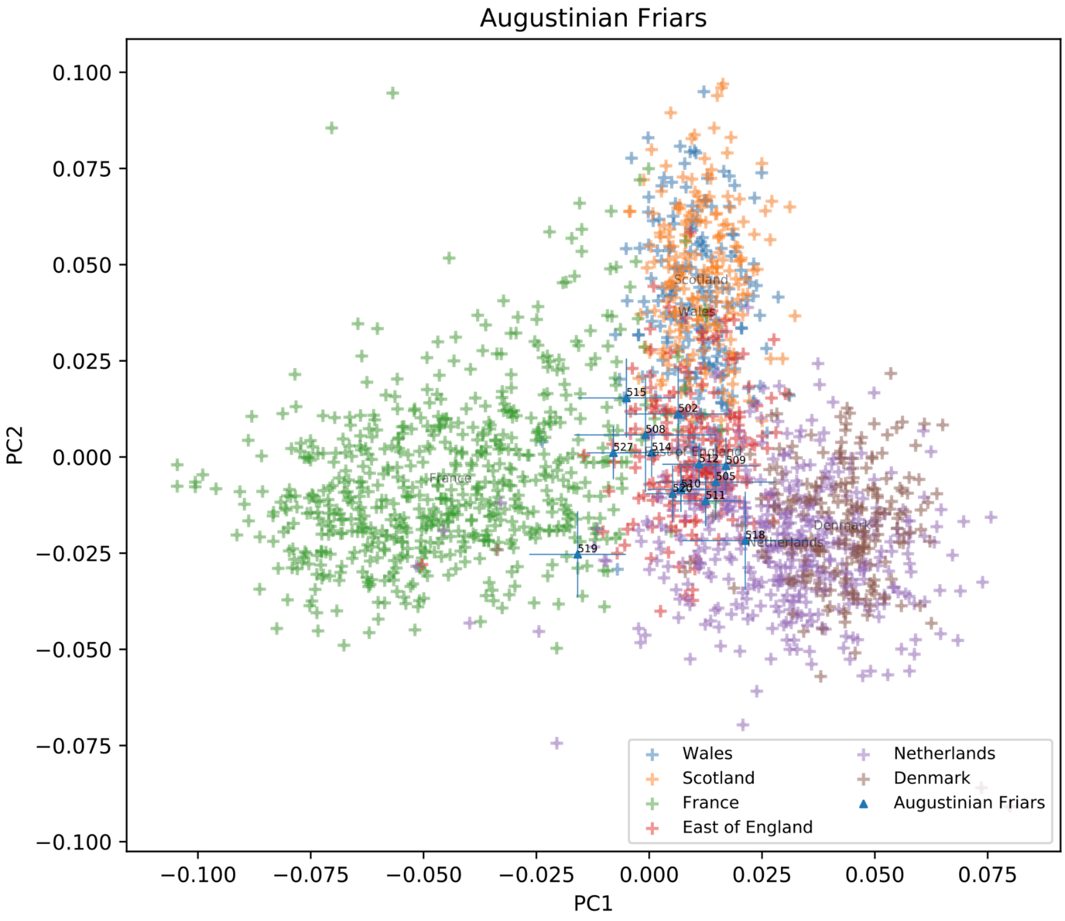


**Figure S6: PCA of West Europe individuals from UK Biobank, showing lsq-projection of historical genomes from Augustinian Friary.**

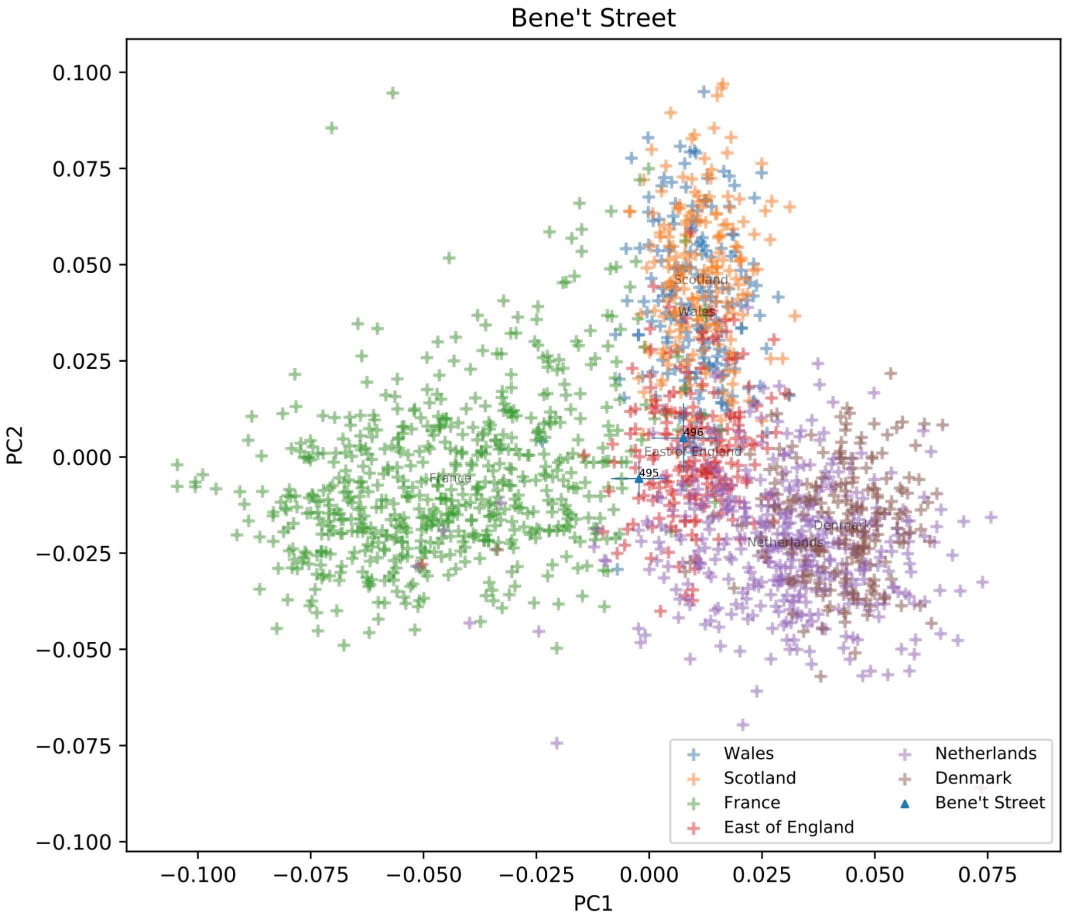


**Figure S7: PCA of West Europe individuals from UK Biobank, showing lsq-projection of historical genomes from Bene’t Street.**

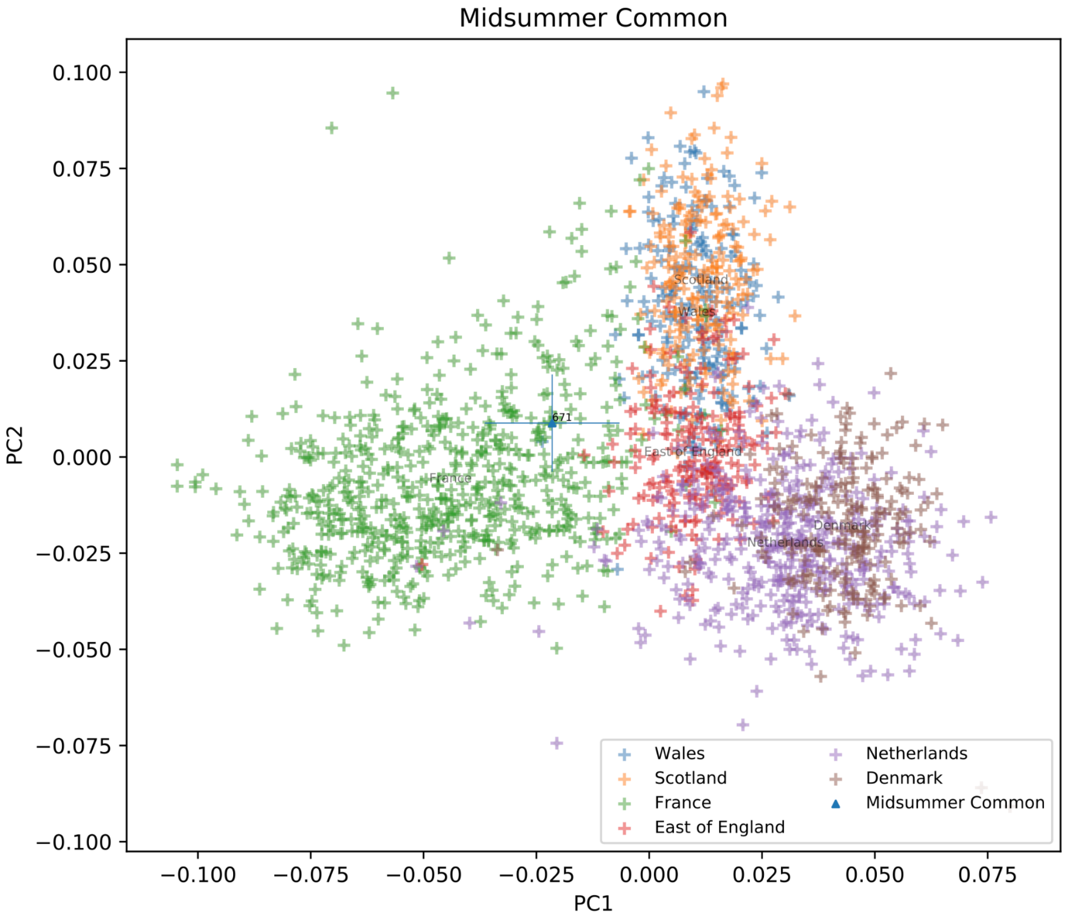


**Figure S8: PCA of West Europe individuals from UK Biobank, showing lsq-projection of historical genomes from Midsummer Common.**

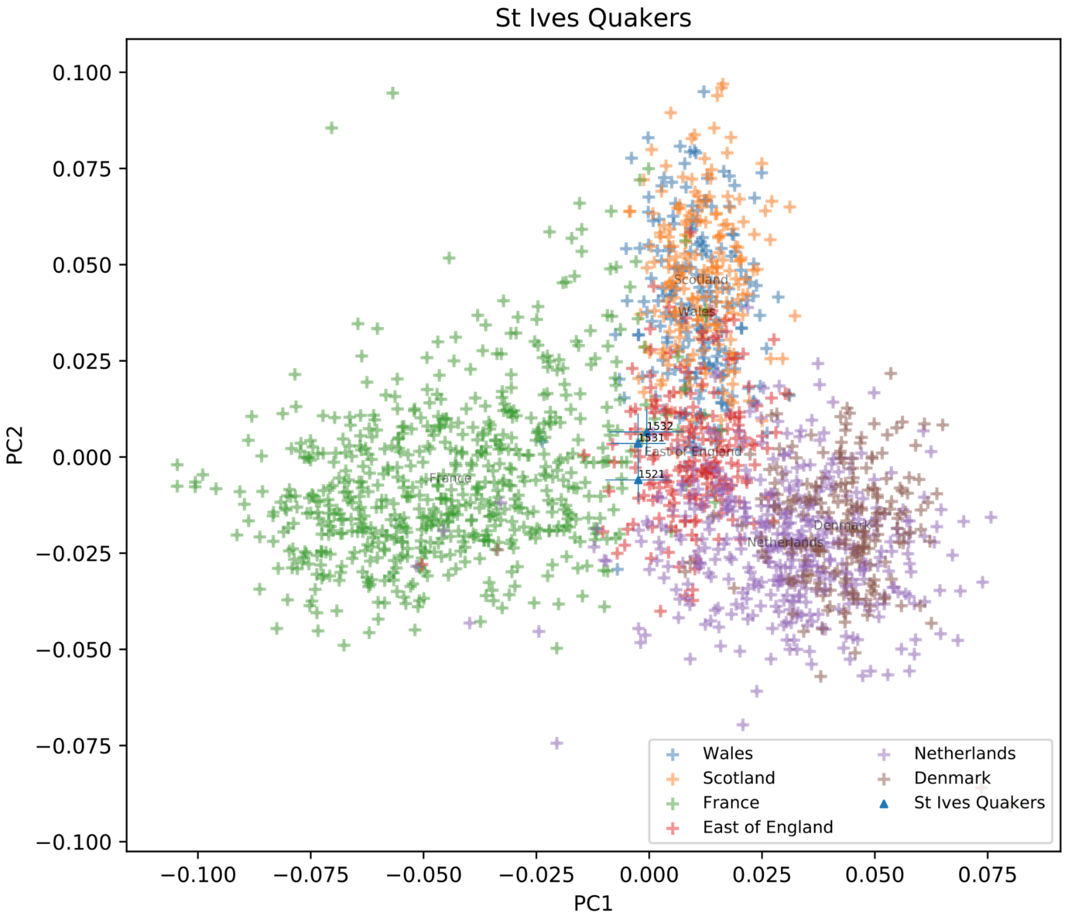


**Figure S9: PCA of West Europe individuals from UK Biobank, showing lsq-projection of historical genomes from Hemingford Gray Quakers.**

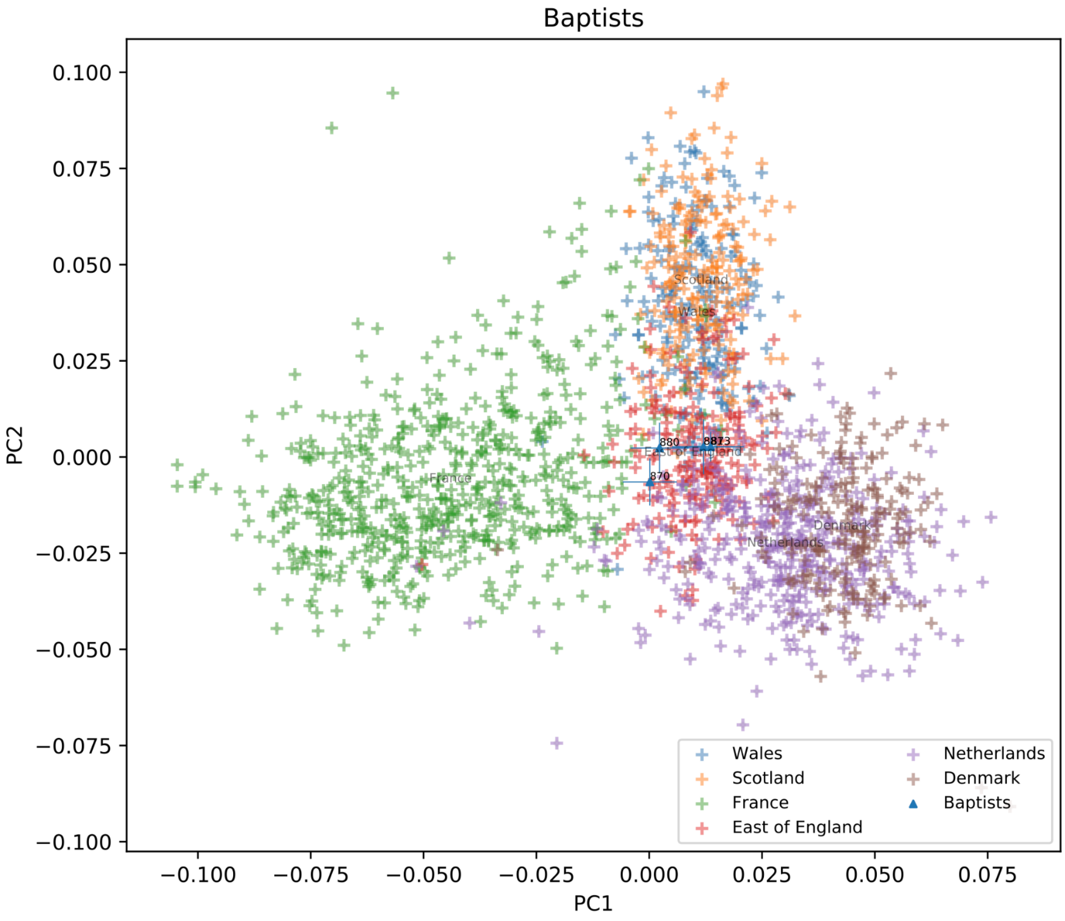


**Figure S10: PCA of west Europe individuals from UK Biobank, showing lsq-projection of historical genomes from Providence Calvinistic Baptist Chapel.**

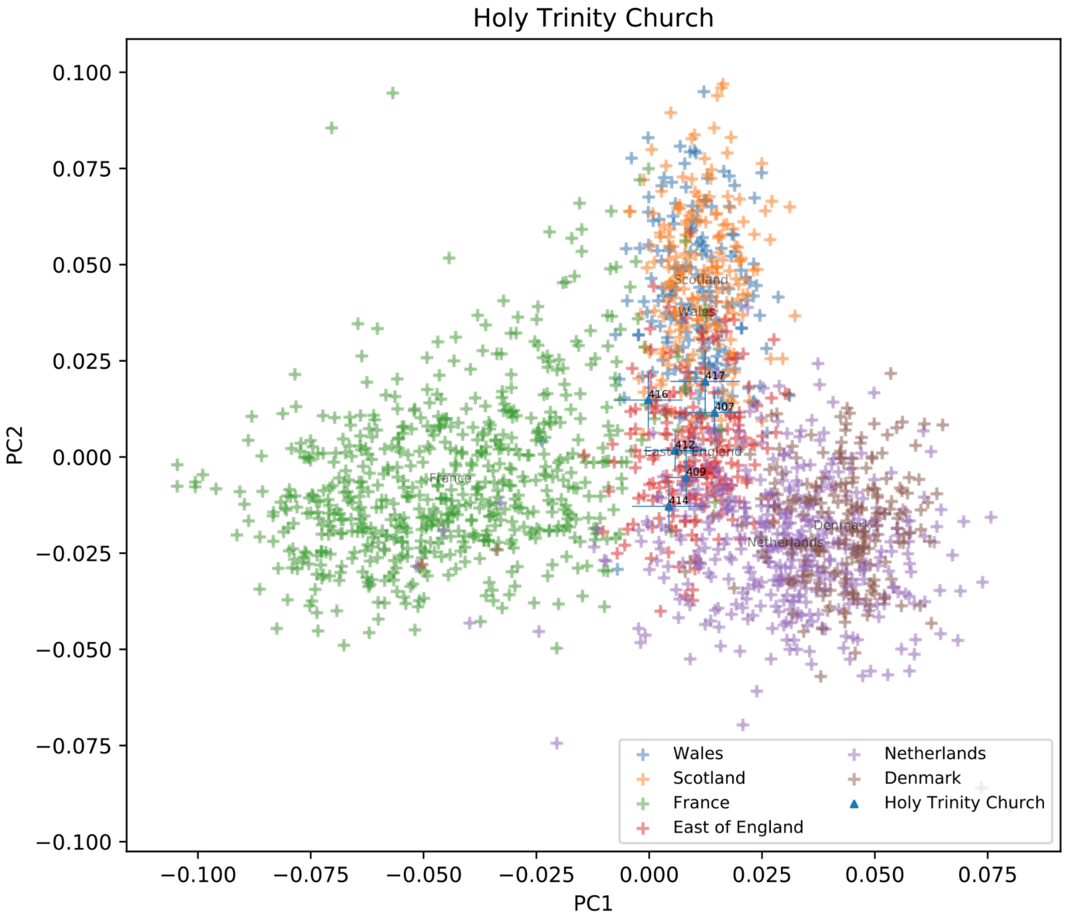


**Figure S11: PCA of west Europe individuals from UK Biobank, showing lsq-projection of historical genomes from Holy Trinity Church.**

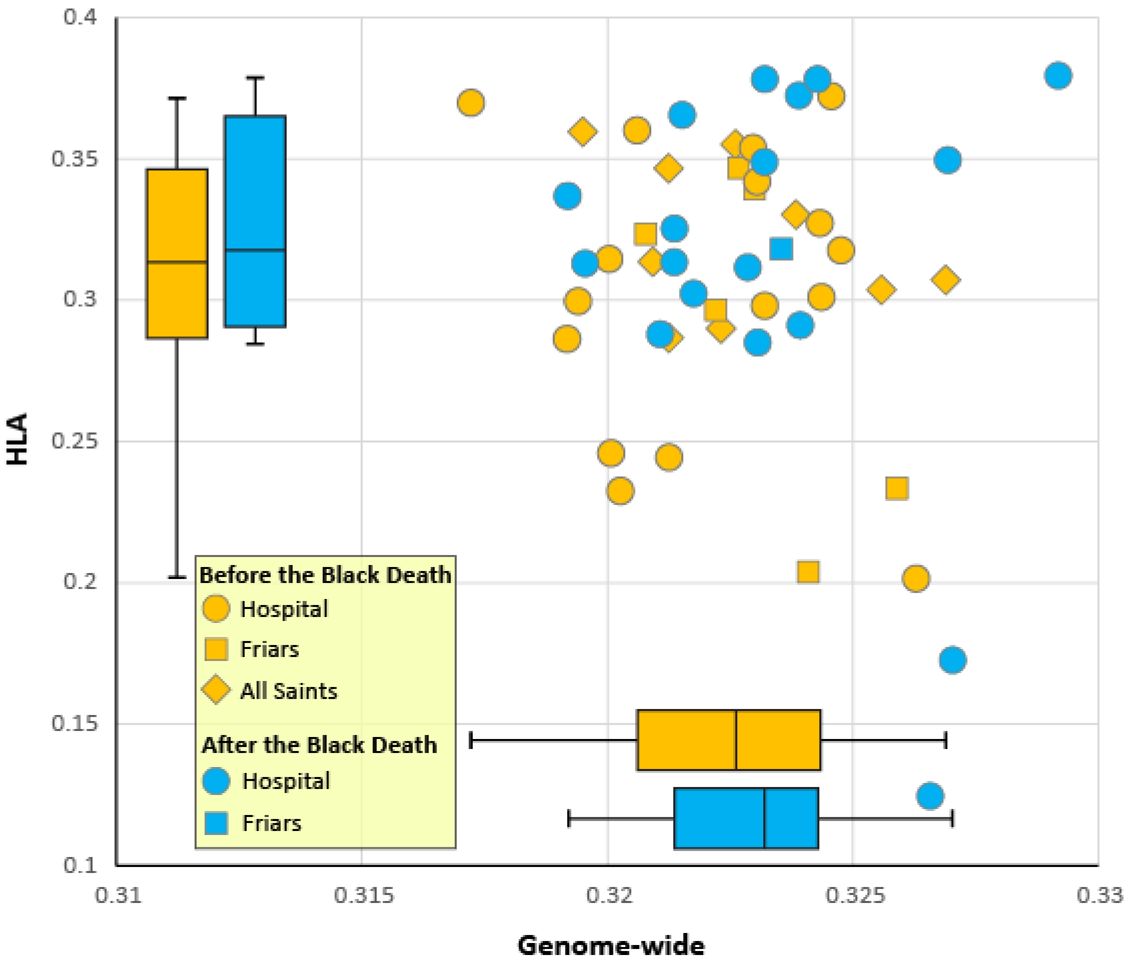


**Figure S12. Heterozygosity of the medieval Cambridge genomes from before and after the Black Death.** Average heterozygosity estimates for 5.4 million variants with MAF>0.05 were obtained from imputed genotypes of 50 genomes with coverage >0.1x from 4 sites in Cambridge.

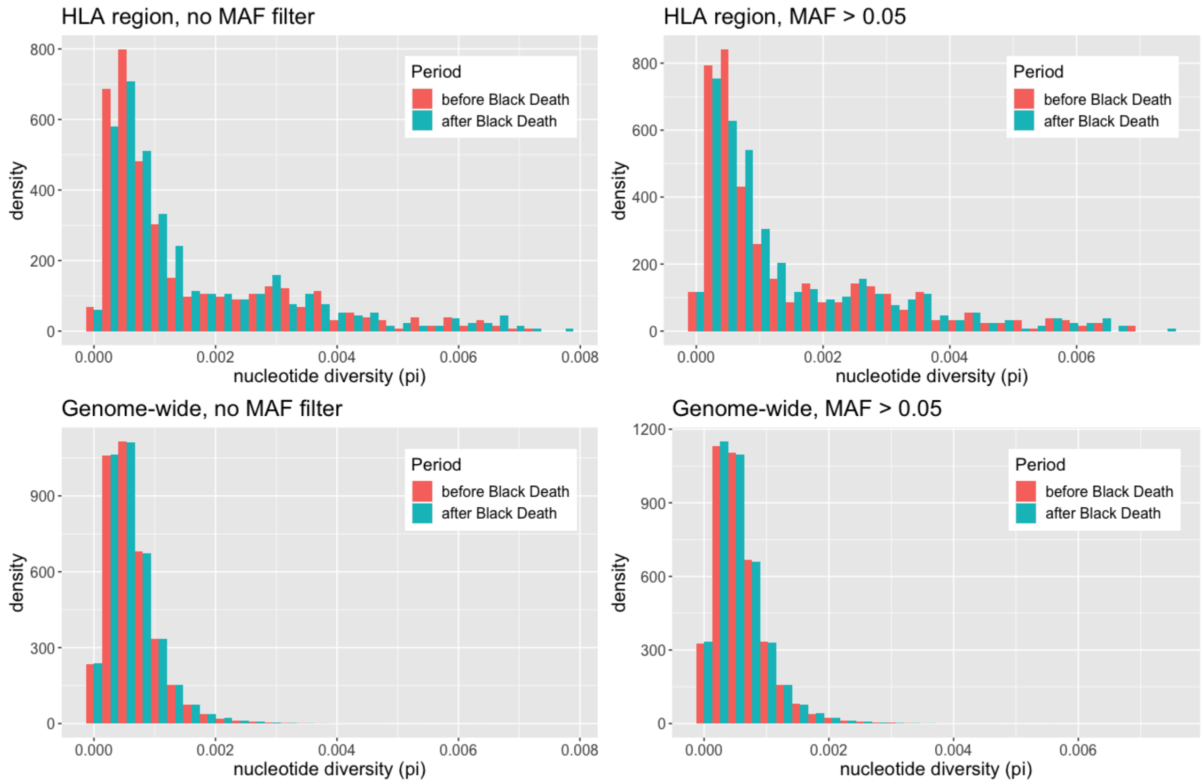


**Figure S13. Histograms comparing nucleotide diversity of the medieval Cambridge genomes from before and after the Black Death.**
